# Molecular identification of wide-field amacrine cells in mouse retina that encode stimulus orientation

**DOI:** 10.1101/2023.12.28.573580

**Authors:** Silvia J. Park, Wanyu Lei, John Pisano, Andrea Orpia, Jacqueline Minehart, Joseph Pottackal, Christin Hanke-Gogokhia, Thomas E. Zapadka, Cheryl Clarkson-Paredes, Anastas Popratiloff, Sarah E. Ross, Joshua H. Singer, Jonathan B. Demb

## Abstract

Visual information processing in the retina is sculpted by a diverse group of inhibitory interneurons called amacrine cells. For most of the >60 amacrine cell types, molecular identities and specialized functional attributes remain unknown. Here, we developed an intersectional genetic strategy to target a group of wide-field amacrine cells (WACs) in mouse retina that co-express the transcription factor Bhlhe22 and the Kappa Opioid Receptor (KOR; B/K WACs). B/K WACs feature straight, unbranched dendrites spanning over 0.5 mm (∼15° visual angle) and produce non-spiking responses to either light increments or decrements. Two-photon dendritic population imaging reveals Ca^2+^ signals tuned to the physical orientations of B/K WAC dendrites, signifying a robust structure-function alignment. B/K WACs establish divergent connections with multiple retinal neurons, including unexpected connections with non-orientation-tuned ganglion cells and bipolar cells. Our work sets the stage for future comprehensive investigations of the most enigmatic group of retinal neurons: WACs.

## INTRODUCTION

Inhibitory interneurons modulate information flow in the brain depending on their diverse morphologies and synaptic mechanisms. Indeed, the remarkable diversity of inhibitory interneurons spans various brain regions, including the cortex, hippocampus, spinal cord, and retina (Booker & Vida, 2018; He et al., 2016; Sweeney et al., 2018; Yan et al., 2020). A comprehensive understanding of the structure and function of inhibitory interneurons is crucial to decipher the functional dynamics and organizational principles of neural circuits.

In the retina, visual information is first encoded by photoreceptors, which transmit signals to the output neurons, the retinal ganglion cells (RGCs), via multiple, parallel excitatory pathways: photoreceptors ◊ bipolar cells ◊ RGCs (Baden et al., 2016; Bae et al., 2018; Euler et al., 2014; Goetz et al., 2022; Kerschensteiner, 2022; Tran et al., 2019). Signaling in these pathways is modulated and shaped by the major class of inhibitory interneurons, known as amacrine cells, which are the most diverse yet least understood retinal cell class (MacNeil & Masland, 1998; Masland, 2012; Yan et al., 2020). Amacrine cells provide inhibitory synaptic input to bipolar cell terminals, RGC dendrites and other amacrine cells (Demb & Singer, 2015; Diamond, 2017).

Classically, amacrine cell function is understood in the context of lateral inhibition that generates RGC receptive field surrounds (Farrow et al., 2013; Flores-Herr et al., 2001; Werblin, 1972; Zaghloul et al., 2007). Amacrine cell functional diversity, however, extends well beyond this classical operation, as demonstrated by specialized computations in various cell types. For example, GABAergic synapses made by starburst amacrine cells are activated preferentially by specific directions of motion (Euler et al., 2002; Jain et al., 2020; Koren et al., 2017), which drives direction-selective (DS) firing in DS-RGCs (Briggman et al., 2011; Lee et al., 2010; Wei, 2018); VGluT3 amacrine cells are sensitive to local motion and provide excitatory input that generates object-motion selectivity in the downstream W3 RGCs (Kim et al., 2015; Krishnaswamy et al., 2015; Lee et al., 2014); S–cone amacrine cells are tuned to blue-ON signals, and their inhibitory output putatively drives blue-OFF responses in downstream RGCs (Chen & Li, 2012; Sher & DeVries, 2012). Therefore, understanding how amacrine cells process visual features is imperative for deciphering retinal computation.

Systematic understanding of the diversity of amacrine cells and their functional roles relies on the ability to identify specific cell types and to interrogate systematically their roles in retinal circuits (Jo et al., 2018; Zhu et al., 2014). Based on dendritic field diameters, amacrine cells can be divided into narrow-field (**∼**30 − 150 μm), medium-field (**∼**150 − 500 μm), and wide-field (>500 μm) subclasses (Masland, 2012). Among these, wide-field amacrine cells (WACs) are the least understood subclass. Stochastic labeling of WACs revealed dendrites that extend more than 0.5 mm from the soma, with a diameter of more than 1 mm (30° of visual angle) in mice, making them the largest cells in the retina (Badea & Nathans, 2004; Lin & Masland, 2006; Perez De Sevilla Muller et al., 2007; Zhu et al., 2014). Systematic explorations of their functions and synaptic connectivity, however, have been hindered due to the lack of genetic approaches for targeted functional recording and manipulation. In this study, we introduce an intersectional genetic approach to target a subset of non-spiking WACs featuring straight, unbranched dendrites. This approach enabled us to investigate both the functional properties and synaptic partnerships of these WACs, laying the foundation for future comprehensive studies of the roles these cells play in retinal information processing.

## RESULTS

### Intersectional genetic approach for studying WACs

Our intersectional genetic approach to identify specific WACs combines three transgenic alleles: Cre and Flpo recombinases driven by two distinct genetic loci, and a reporter of dual (Cre- and Flpo-mediated) recombination (Cai, Kardon, et al., 2016; Jo et al., 2018) (**Figure 1A**). In our search for markers of WACs, the transcription factor Bhlhe22 (also known as Bhlhb5) stood out. Bhlhe22 has an essential role in the development of a spectrum of GABAergic amacrine cells as well as type 2 OFF cone bipolar cells (Feng et al., 2006; Huang et al., 2014). Indeed, an existing transcriptomic database suggested that approximately half of GABAergic amacrine cells express mRNAs encoding *Bhlhe22* at a relatively high level (Yan et al., 2020) (**Figure 1B**).

**Figure 1.**
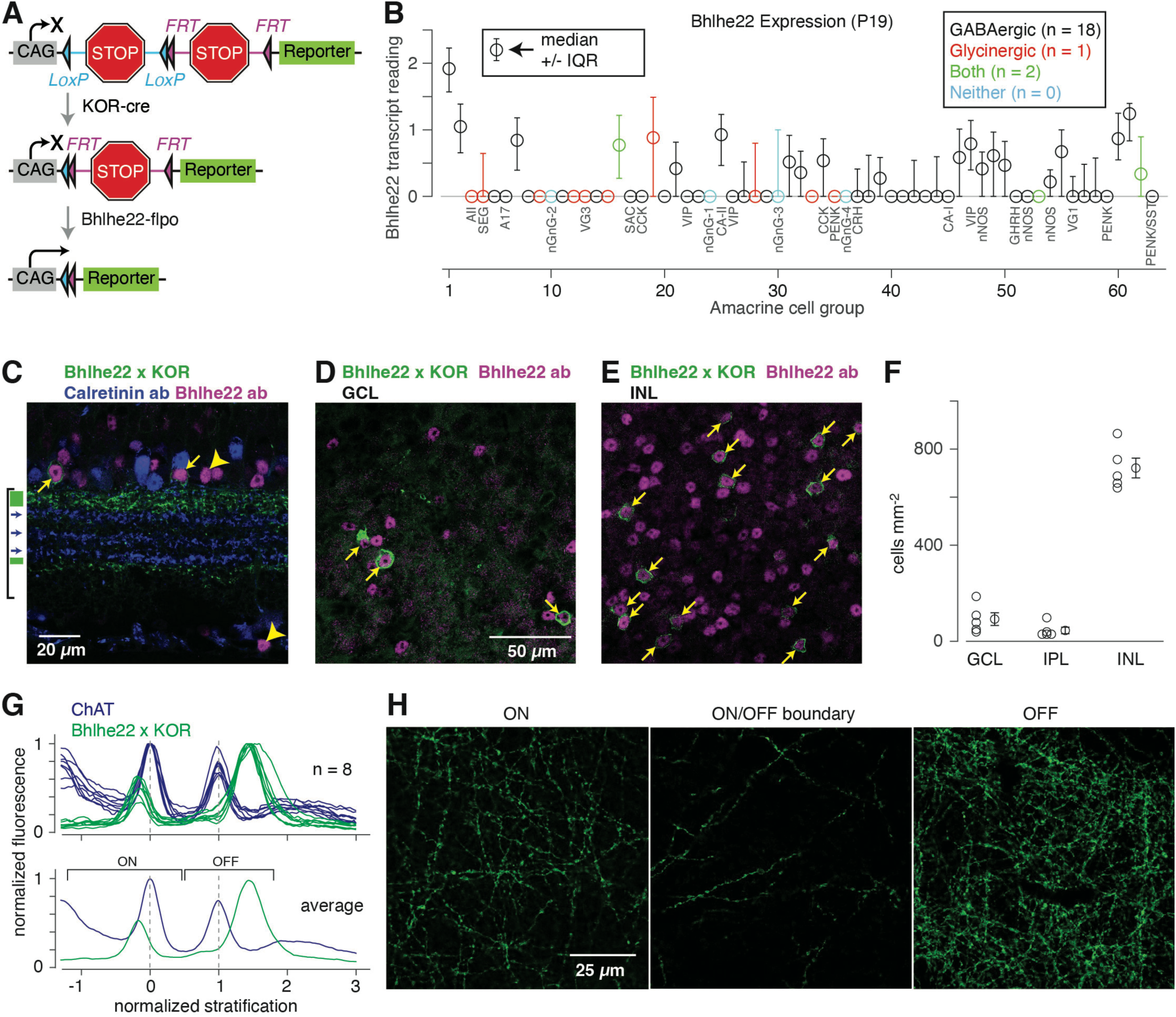
Genetic strategies target *Bhlhe22*-expressing amacrine cells and cells that co-express *Bhlhe22* and KOR (B/K). **A.** Intersectional genetic strategy. A KOR-Cre allele, acting on Loxp sites, combined with the *Bhlhe22*-Flpo allele, acting on Frt sites, removes two stop sequences and induces the expression of a reporter gene (either ReaChR-mCitrine fusion protein or tdTomato), driven by the CAG promoter. **B.** *Bhlhe22* transcript readings in previously identified groups of amacrine cells (Yan et al., 2020). Data points show the median *Bhlhe22* transcript reading and the inter-quartile range (IQR) of these readings within each group. Cell groups are identified as GABAergic, glycinergic, both or neither, according to the definitions in the original study, and groups that correspond to known cell types are labeled with the name of the type (e.g., AII or A17 amacrine cell) or a characteristic gene name (e.g., VIP^+^ or nNOS^+^ amacrine cells). **C.** Intersection between KOR-Cre and *Bhlhe22*-Flpo labels amacrine cell processes primarily in two layers (green rectangles, left) relative to calretinin bands (blue arrows; bracket marks the IPL). Top and bottom calretinin bands are identical to the OFF and ON ChAT bands, respectively. B/K amacrine cells were labeled with Bhlhe22 antibody (yellow arrows); the antibody also labeled amacrine cells without reporter expression (yellow arrowheads). **D.** Whole-mount retina showing B/K amacrine cells in the GCL with Bhlhe22 antibody labeling (arrows). **E.** Same as D. for the INL, which shows a higher density of labeled B/K somas. **F.** Density of B/K cells in different layers (n = 5 retinas) based on counts within 0.102 mm^2^ regions. Error bars indicate ± SEM across samples. **G.** Fluorescence profile of B/K intersectional genetic labeling relative to the ChAT bands in eight individual z-stacks (top). The average (bottom) shows fluorescence relative to the layers where ON and OFF bipolar cells stratify their axon terminals. **H.** Example images of long, straight B/K amacrine cell processes in ON and OFF layers of the IPL and the boundary between these layers.

Given that WACs are primarily GABAergic (Wassle & Boycott, 1991; Zhu et al., 2014), whereas narrow-field cells are primarily glycinergic (Wassle et al., 2009), it is likely that unidentified Bhlhe22-expressing (*Bhlhe22*^+^) cells include uncharacterized WAC types. Indeed, the *Bhlhe22*-Flpo transgenic line, crossed with a Flpo-dependent reporter, labeled type 2 bipolar cells as well as amacrine cell processes that stratified at four levels of the inner plexiform layer (IPL) (Cai, Kardon, et al., 2016); and our pilot studies suggested that labeled amacrine cells include WACs.

To increase the genetic specificity in targeting WACs, we crossed the *Bhlhe22*-Flpo line with several Cre lines and discovered a compelling intersection with the Kappa Opioid Receptor (KOR)-Cre line, which labels amacrine cell processes that stratify primarily in two layers, including a subset of those labeled in the *Bhlhe22*-Flpo line (Cai, Huang, et al., 2016). Indeed, the *Bhlhe22*-Flpo;KOR-Cre intersection, combined with a dual ReaChR-mCitrine reporter line (Hooks et al., 2015) (**Figure 1A**), labeled a group of amacrine cells with processes that stratified primarily at the borders of the IPL (**Figure 1C**) and very rarely labeled bipolar cells or RGCs. Furthermore, nearly all amacrine cells expressing the reporter (98%) also were labeled by Bhlhe22 immunofluorescence (144/147 cells; n = 4 retinas) (**Figure 1C-E**). For simplicity, we refer to the cells labeled by this intersectional genetic strategy as B/K amacrine cells. Notably, we could not definitively match B/K amacrine cells to a subset of *Bhlhe22^+^*cells in the transcriptomic dataset due to the low levels of the *Oprk1* transcript (Yan et al., 2020) (**Figure 1-figure supplement 1**).

Most (>80%) B/K cells had somas in the inner nuclear layer (INL), although some localized to either the ganglion cell layer (GCL) or the inner plexiform layer (IPL; **Figure 1F**). The fluorescence profile of B/K processes, analyzed in stacks of confocal images, peaked in either ON or OFF layers, as defined by the known locations of ON or OFF bipolar cell axon terminals (Borghuis et al., 2013; Franke et al., 2017) (**Figure 1G**). Confocal images showed long, straight processes concentrated near the fluorescence peaks in ON or OFF regions of the IPL with sparse labeling in between, near the ON/OFF boundary (**Figure 1H**).

### B/K amacrine cells have long, unbranched processes with oriented dendritic trees

To investigate the morphology and functional properties of single B/K amacrine cells, a cell in either the GCL or INL (but not the IPL) was targeted for whole-cell recording (Park et al., 2015; Park et al., 2020; Park et al., 2018). The targeted cell was filled with Lucifer Yellow through the pipette, and, after fixation, the dendrites were imaged by confocal microscopy (**Figure 2A, B**). Dendrite stratification was normalized to the stratification of the ON and OFF ChAT (choline acetyltransferase) bands, which correspond to two sets of cholinergic starburst amacrine cell dendrites (**Figure 2C**) (Manookin et al., 2008; Park et al., 2015; Park et al., 2020; Park et al., 2018). Low-resolution images of B/K WACs were traced to determine the number of extended dendrites, after initial branching within ∼50 µm of the soma (**Figure 2A, B**). In many cases, the entire dendrite could not be captured because the Lucifer Yellow signal faded or because of cuts in the tissue. Nevertheless, dendrites were commonly >0.5 mm in length, consistent with the general WAC morphology. Dendrites of ON cells primarily stratified on the proximal side of the ON ChAT band, and those of OFF cells primarily stratified on the distal side of the OFF ChAT band; although four cells in the sample (1 ON, 3 OFF) stratified between the bands (**Figure 2D**). For the entire sample (n = 55 cells), dendrite number versus stratification level showed two prominent clusters: OFF cells with ∼4-6 dendrites and ON cells with ∼7-13 dendrites (**Figure 2D**). Stratification was measured within ∼70-100 µm of the soma, but the dendrite retained its initial stratification level in distal dendrites, ∼500 µm from the soma (**Figure 2E**). Overall, B/K amacrine cells apparently comprise several WAC types with different stratification levels.

**Figure 2.**
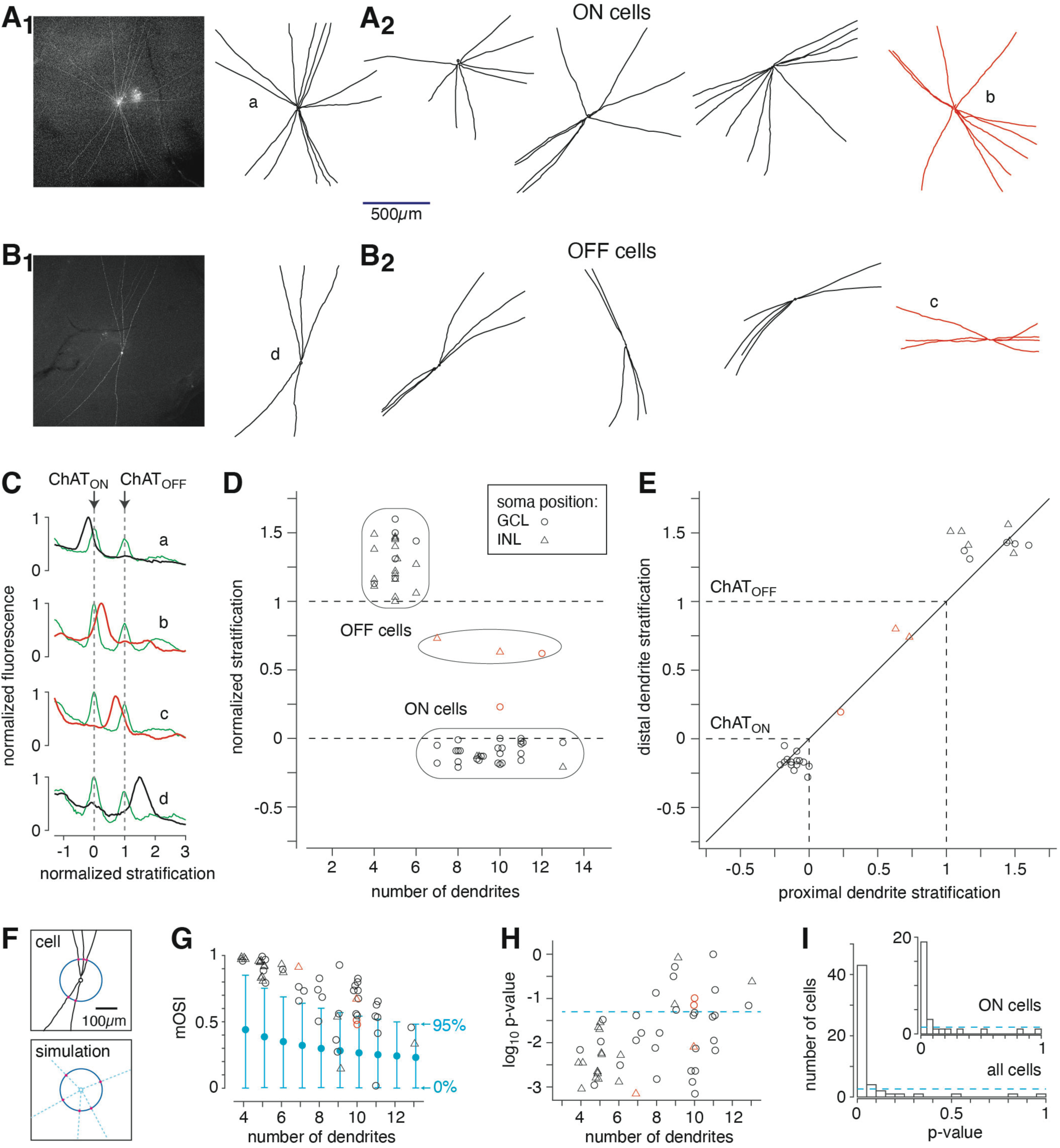
B/K amacrine cells have long, unbranched dendrites and stratify at distinct levels of the inner plexiform layer. **A_1._** Confocal image (left) and tracing (right) of a B/K wide-field amacrine cell (WAC) in the ON layer. Scale bar in A applies to all cells in A and B. **A_2_.** Four additional ON B/K WACs. Cell at far right stratified between the ChAT bands. **B_1_, B_2_.** Same as A for OFF B/K WACs. Cells labeled by lower-case letters in A and B correspond to the individual cells whose stratification is shown in C. The red cells in A and B stratify between the ChAT bands, as shown in C. **C.** Fluorescence profiles of labeled dendrites of example ON and OFF WACs in A and B. Stratification is shown for each of four individual cells relative to ChAT bands labeled in the same tissue. Stratification is normalized to the inner/ON ChAT band (0, ChAT_ON_) and the outer/OFF ChAT band (1, ChAT_OFF_). **D.** Population analysis (n = 55 cells) showing the number of dendrites relative to the stratification, normalized to the ChAT bands. Apparent groups of cells are shown with outlines. Soma position is indicated by the symbol. **E.** A subset of B/K WAC dendrites were visualized near the soma (within ∼150 µm) and at a distal location (∼500 µm from the soma). Dendrites remained in their original layer, as indicated by the points falling near the identity line. Symbols indicate soma location, as in D. **F.** Top, example OFF B/K WAC showing dendrite angle (red dots on circle with 100 µm radius) relative to soma. Bottom, simulated cell from bootstrapped distribution with the same number of dendrites. **G.** The morphological orientation selectivity index (mOSI) for all B/K WACs (symbols match those in D.). Monte Carlo null distributions of mOSI values for each number of dendrites are shown in cyan; the upper bound indicates the 95^th^ percentile of the null distribution. **H.** The log_10_ p-value of each B/K WAC from G. as a function of dendrite number. Dashed line indicates p = 0.05. **I.** Distribution of p-values for all B/K WACs relative to a uniform distribution (dashed line), the null hypothesis if there were no orientation bias of the dendritic trees. ON B/K WACs, analyzed separately, are shown in the inset.

Most of the B/K WACs appeared to have asymmetric, oriented dendritic trees. This was especially true for the OFF cells. To test for an orientation bias, we quantified dendrite angles at the point they crossed a circle (radius = 100 µm) centered on the soma **(Figure 2F)**. We computed a morphological orientation selectivity index (mOSI), based on a previous OSI used to quantify orientation selectivity of physiological responses, which were characterized by a response amplitude at each of multiple stimulus orientations (Liang et al., 2023; Niell & Stryker, 2008). Here, the mOSI value reflected the set of dendrite angles for each cell, and the amplitude value for each dendrite was assigned a value of 1.0 (i.e., we did not account for dendrite length; see Methods). Each observed mOSI value was compared to a Monte Carlo null distribution of mOSI values for the same number of dendrites (4-13 per cell; see Methods) and was assigned a p-value (see Methods). Most B/K WACs had mOSI values that exceeded the 95^th^ percentile of the null distribution (**Figure 2G**). Indeed, all OFF B/K WACs (i.e., those with 4-6 dendrites) had significant mOSIs (p < 0.05) individually, and about half of the ON B/K WACs (i.e., those with 7-13 dendrites) did as well (**Figure 2H**).

To test for orientation bias at the population level, we compared the observed distribution of mOSI p-values to the null hypothesis, which corresponds to a uniform distribution. The distribution of p-values from all B/K WACs was significantly different from a uniform distribution (**Figure 2G**), and this was also true for the ON cells analyzed separately (**Figure 2G**, inset). We conclude that B/K WACs show oriented dendritic trees as a population, although some individual ON cells show relatively symmetrical dendrites. We did not perfectly align the retinas to the body axis, and so we could not perform a quantitative analysis of relative dendrite angle in this space. However, we presume that the B/K WAC population comprises all angles of dendrite orientation.

### B/K amacrine cells do not fire conventional action potentials

The stratification of amacrine cell dendrites relative to the terminals of ON and OFF bipolar cells in most cases predicts either ON or OFF responses to contrast modulation (Euler et al., 2014)(for exceptions, see (Jo et al., 2023; Park et al., 2020)). Indeed, B/K WACs with dendrites in the OFF layer depolarized during the negative/OFF phase of a contrast-reversing spot stimulus and hyperpolarized during the positive/ON phase, whereas B/K WACs with dendrites in the ON layer depolarized slightly during the ON phase and hyperpolarized strongly during the OFF phase (**Figure 3A**). For both ON and OFF B/K WACs, responses to increasing spot diameters showed increased modulation amplitude over a distance of ∼1 mm (**Figure 3B**), consistent with the diameter of their dendritic trees.

**Figure 3.**
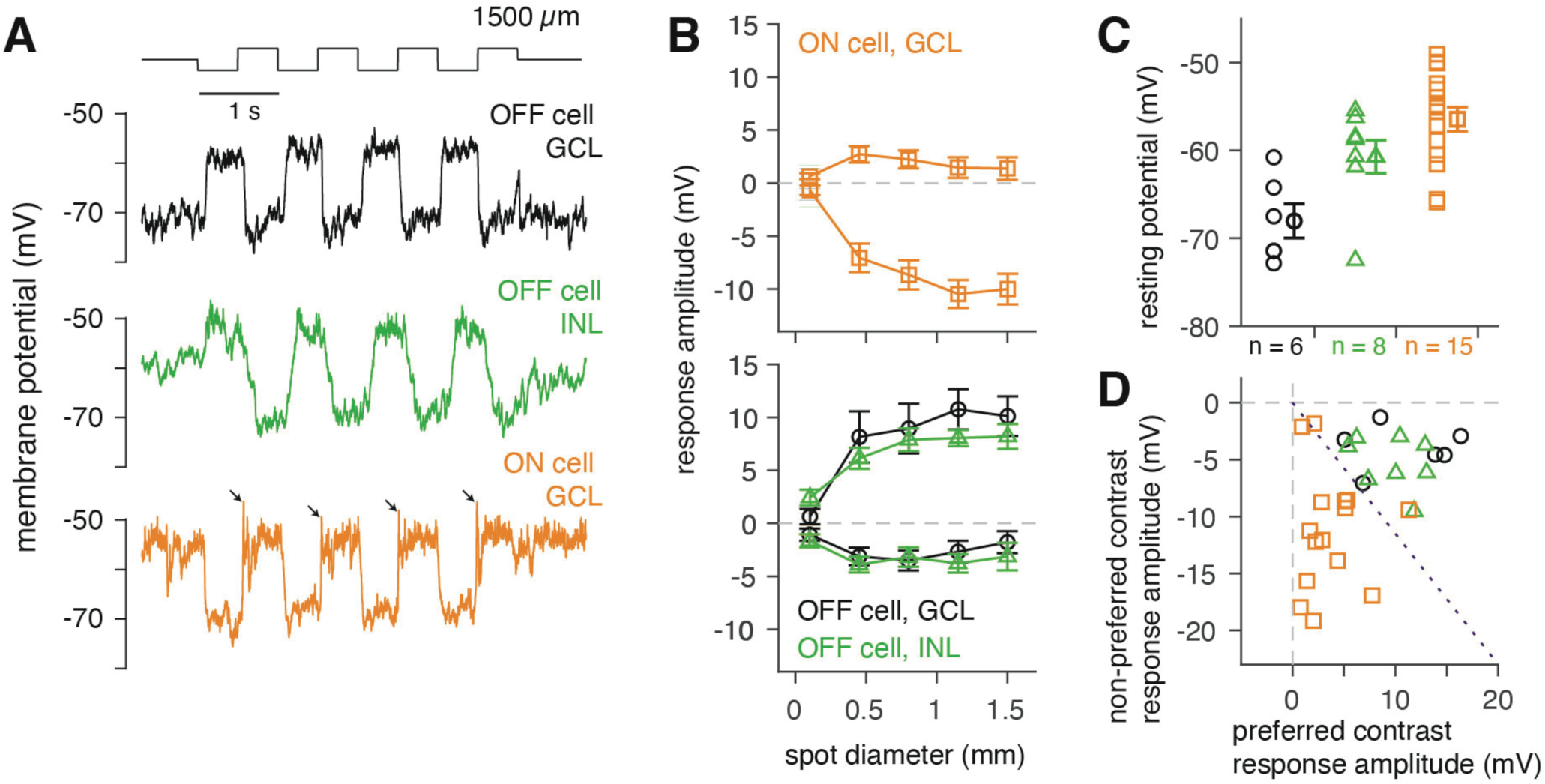
B/K amacrine cells comprise ON and OFF types and do not fire action potentials. **A.** Membrane potential recording of response to a contrast-reversing spot stimulus (1.5-mm diameter) for three B/K WACs, including two OFF cells with somas in either the GCL or INL and an ON cell with a soma in the GCL. For the ON cell, small fast events were observed during the initial depolarizing response (arrows). **B.** Responses to contrast-modulated spots of varying diameters. Response amplitudes to the dark (OFF) and bright (ON) phases of the spot are quantified relative to the resting potential and distinguish ON and OFF B/K WACs: ON WACs depolarize modestly to bright stimuli and hyperpolarize strongly to dark, whereas OFF WACs show the opposite pattern. Error bars in G. and I. indicate ± SEM across cells. The primary response (depolarization for OFF cells, hyperpolarization for ON cells) asymptotes at ∼1 mm diameter. **C.** Resting membrane potential of B/K WACs in the presence of a mean luminance. OFF B/K WACs in the GCL were more hyperpolarized than OFF B/K WACs in the INL (t(12) = 2.66, p = 0.02) or ON B/K cells in the GCL (t(19) = 4.64, p < 0.001), whereas OFF B/K cells in the INL and ON B/K cells in the GCL were more similar (t(21) = 1.84, p = 0.08, two-sample t-tests). **D.** Peak depolarization to the preferred stimulus versus peak hyperpolarization to the non-preferred stimulus. Cells with equal modulation around the resting potential would fall along the dashed line. The combined group of OFF B/K WACs had larger depolarizing responses to preferred contrast compared to ON B/K WACs (t(27) = 5.29, p < 0.001), whereas ON B/K WACs had relatively larger hyperpolarizing responses to non-preferred contrast (t(27) = 4.63, p < 0.001).

We did observe some heterogeneity in the B/K cells: the OFF B/K cells with somas in the GCL had relatively hyperpolarized resting potentials compared to the other groups (**Figure 3C**); and OFF B/K WACs in the INL and GCL had similar patterns of response to contrast modulation, with relatively large depolarizing responses to preferred contrast, whereas ON B/K cells had relatively large hyperpolarizing responses to non-preferred contrast (**Figure 3D**). Notably, none of the recorded cells fired conventional action potentials (spikes). This differs from the response of another class of spiking WACs, which have distinct axonal and dendritic compartments (e.g., nNOS-1 and dopaminergic amacrine cells), including WACs recorded under the same conditions as those presented here (Badea & Nathans, 2004; Greschner et al., 2014; Kim et al., 2022; Lin & Masland, 2006; Murphy-Baum & Taylor, 2015; Newkirk et al., 2013; Park et al., 2020; Park et al., 2018; Stafford & Dacey, 1997; Taylor, 1996). Nonetheless, some cells exhibited small fast events (e.g., the ON cell in **Figure 3A**) that could reflect voltage-gated channel activity within the dendrites (Bloomfield & Volgyi, 2007), but this was not investigated further.

### B/K WACs exhibit orientation-selective dendritic Ca^2+^ signals

The non-spiking nature of the B/K WACs suggests that they encode information through their dendrites. Further, their unique, highly oriented dendritic architecture evokes parallels with the elongated dendritic fields of a variety of orientation-selective (OS) and direction-selective (DS) neurons (Antinucci et al., 2016; Bloomfield, 1994; Kim et al., 2008; Kim et al., 2014; Nath & Schwartz, 2017; Weiler et al., 2022). To test whether B/K WACs encode orientation or direction information within their dendrites, we expressed the Ca²⁺ sensor GCaMP7s in B/K WACs using a recently developed dual reporter line (Ai195) (Qi et al., 2024) and performed population dendritic two-photon imaging at multiple depths throughout the IPL while presenting gratings drifting in eight directions of motion. In both the OFF (**Figure 4-video 1**) and ON layers (**Figure 4-video 2**), B/K dendrites exhibited substantial responses for two opposite directions of motion that shared the same orientation (**Figure 4A**). For individual regions of interest (ROIs), we computed both orientation- and direction-selective indices (OSI, DSI; see Methods). For the vast majority of ROIs, the OSI value was significantly larger than the DSI value (**Figure 4B**). ROIs with large OSI values were found at multiple levels of the IPL (**Figure 4C**).

**Figure 4.**
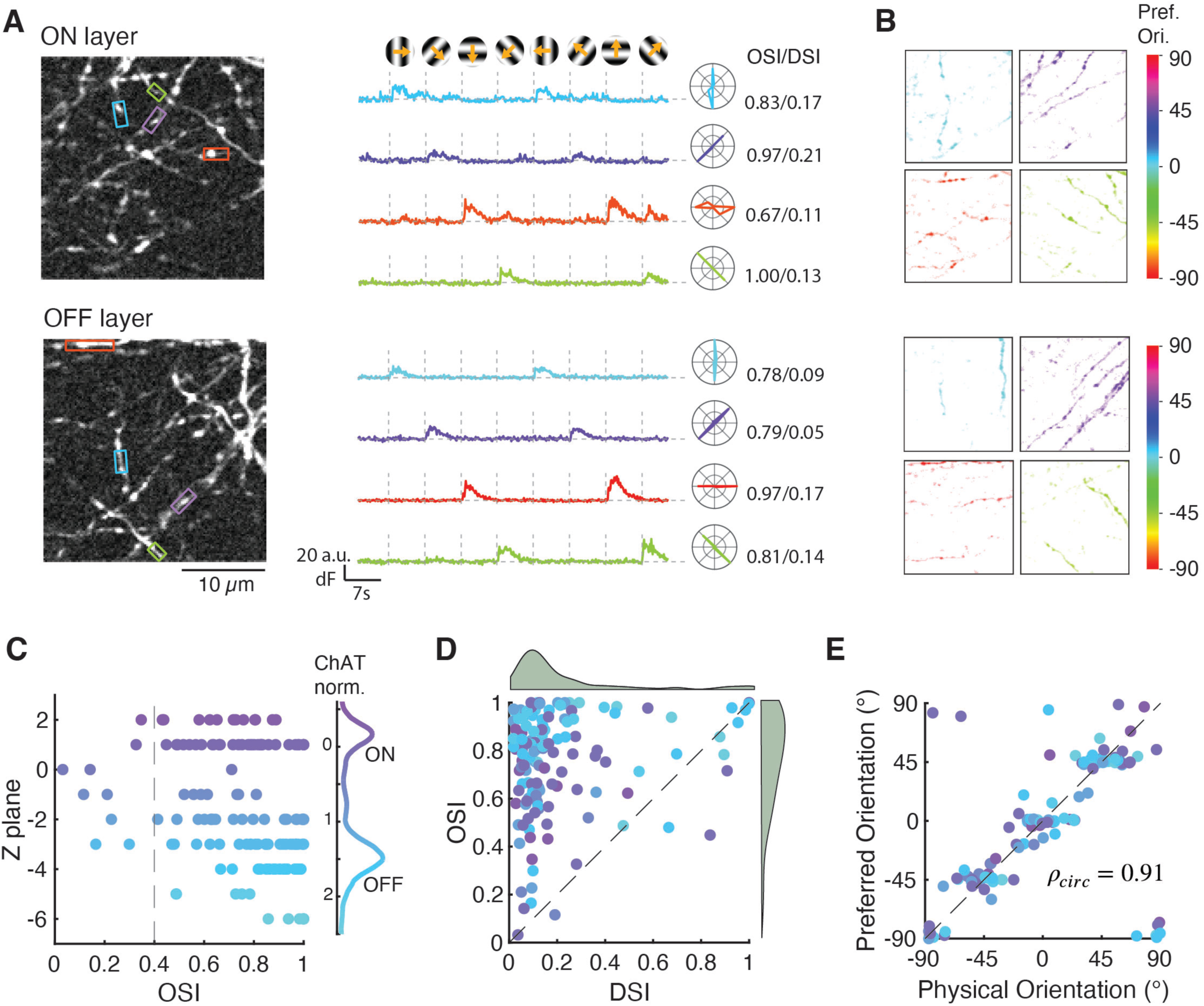
Population dendritic imaging reveals orientation-selective B/K dendrites at different layers in the inner plexiform layer (IPL). **A.** Example imaging planes for ON (top) and OFF (bottom) layers. Each plane shows a local correlation images over time. Example regions of interest (ROIs) are rectangles whose long axis aligns with the orientation of the dendrite. Corresponding baseline-subtracted fluorescence traces (in arbitrary units, a.u.) and polar plots of orientation tuning are shown on the right. Drifting grating stimuli were sinusoidal gratings moving in eight directions. Each stimulus was presented for 2 s, starting at a vertical dashed line, with a 5 s interval between stimuli. The grating had a 2-Hz temporal frequency, a 0.139 cycles/degree (4.54 cycles/mm) spatial frequency and was presented in a patch of 880 µm (29.3 degrees) in diameter. **B.** Spatial footprints tuned to four orientations extracted by sparse non-negative matrix decomposition for the exemplary ON (top) and OFF (bottom) layers shown in A. **C.** Orientation selectivity index (OSI) of dendritic ROIs (n = 135) at different IPL depths. **D.** OSI versus Direction-selective index (DSI) of B/K dendrites at different depths of the IPL. The smoothed probability densities/kernel densities of ROI OSI and DSI are plotted at the margins. OSI (0.75 ± 0.02; n = 135) is significantly larger than DSI (0.22 ± 0.02; t(134) = 20.91, p < 0.001). **E.** Physical orientation versus preferred orientation of B/K dendrite segments with OSI > 0.4 at different depths of the IPL (n = 126 ROIs). The circular Pearson correlation between physical and preferred orientations is 0.91 (p < 0.001). Apparent clustering of orientation preference reflects the limited number of orientations (four) presented combined with the narrow orientation tuning of the ROIs.

Next, we evaluated the relationship between the preferred orientation tuning and the physical orientation of a dendrite. First, to visualize dendrites with common response properties from their overlapping neighbors, we utilized sparse non-negative matrix factorization (sNMF) for spatial footprint extraction from the time-lapse movies (Pnevmatikakis et al., 2016; Wang & Zhang, 2013). This analysis revealed groups of straight, unbranched processes that resemble dendritic segments of filled B/K cells (**Figure 2A**). Each group of processes exhibited a functional orientation preference that closely aligned with their physical orientations (**Figure 4B**). We next compared the physical orientation to the functional tuning of orientation-selective ROIs (i.e., those with OSI > 0.4) and found a strong structure-function correlation (**Figure 4E**) (Lei et al., 2024).

Collectively, we showed that B/K WAC dendrites encoded stimulus orientations that parallel their physical orientations. In theory, the orientation-selective Ca²⁺ responses could further drive orientation-tuned GABA release onto postsynaptic neurons.

### B/K WACs make synapses with non-orientation tuned ganglion cells

B/K WAC dendrites stratify primarily at levels of the IPL that overlap the dendrites of commonly studied wide-field RGCs that do not encode orientation. Indeed, the OFF Delta cell (also known as OFF-Sustained Alpha RGC) (Krieger et al., 2017; Margolis & Detwiler, 2007; Tagawa et al., 1999) stratifies between the OFF ChAT band and the INL, overlapping the primary stratification of B/K WAC dendrites in the OFF layer (**Figure 5A, B**). The ON Alpha cell (also known as ON-Sustained Alpha RGC or the M4 melanopsin RGC) (Estevez et al., 2012; Krieger et al., 2017) stratifies on the vitreal side of the ON ChAT band, overlapping the primary stratification of B/K WAC dendrites in the ON layer (**Figure 5E, F**). We targeted these RGC types because they could be identified by their large soma size and characteristic light response (Krieger et al., 2017), and we tested whether they receive synapses from B/K WACs using optogenetics.

**Figure 5.**
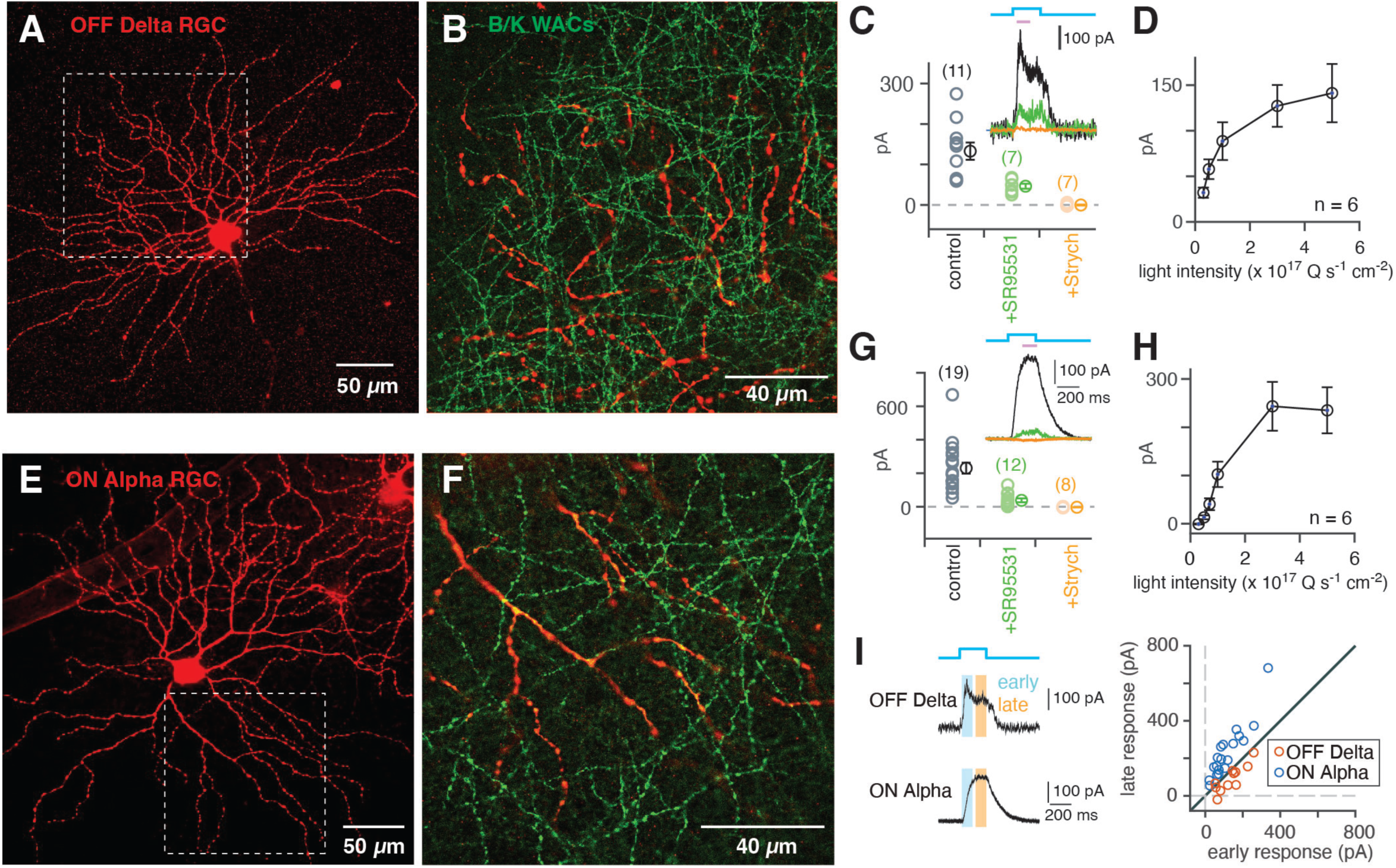
B/K amacrine cells make synapses with non-orientation tuned alpha/delta-type RGCs. **A.** Dendritic tree of a recorded OFF Delta RGC (also known as a Sustained OFF Alpha RGC). Image shows average fluorescence in a confocal stack. Dashed square shows region in B. The RGC was filled with Lucifer Yellow (LY) during whole-cell recording, which was subsequently amplified with LY primary antibody and a red secondary antibody. **B.** Single confocal section showing OFF Delta RGC dendrites from A. relative to B/K WAC dendrites, labeled by the Cre/Flpo-dependent ReaChR-mCitrine reporter. **C.** Optogenetic stimulation (5.3 x 10^17^ Q s^-1^ cm^-2^) of ReaChR-expressing B/K WACs caused an inhibitory postsynaptic current (IPSC, black) in an OFF Delta RGC. The IPSC (133 ± 46 pA; n = 11) was mostly blocked by gabazine (50 µM SR95531; 46 ± 6 pA; n = 7; difference of 106 ± 25 pA, t(6) = 4.22, p = 0.0056) and completely blocked by subsequent addition of strychnine (1 µM; -0.33 ± 1.17 pA; n = 7; difference of 46 ± 5 pA, t(6) = 8.68, p < 0.001). Error bars in all panels indicate ± SEM across cells. Optogenetic experiments were performed in the presence of glutamate receptor blockers (see Methods) to prevent photoreceptor contributions to the light response. Peak IPSC was measured (mauve bar). **D.** Increasing optogenetic stimulation caused increasing IPSC amplitude in OFF Delta RGCs. **E-H.** Same as A-D. for ON Alpha RGCs (also known as Sustained ON Alpha RGCs). The ON Alpha RGC IPSC (231 ± 32 pA; n = 19) was mostly blocked by gabazine (38 ± 12 pA; n = 12; difference of 214 ± 41 pA, t(11) = 5.23 p < 0.001) and completely blocked by strychnine (−2.9 ± 0.8 pA, n = 8; difference of 58 ± 14 pA, t(7) = 4.24; p = 0.0038). **I.** Left, the early (20 – 120 ms) and late (150 – 250 ms) windows of the optogenetic response. Right, plot of early and late responses for each cell. ON Alpha cells had a lower early:late response ratio (0.491 ± 0.033) compared to OFF Delta cells (1.62 ± 0.24; one outlier removed) (t(27) = 6.39; p < 0.001).

After blocking transmission from photoreceptors (Park et al., 2015; Park et al., 2020; Park et al., 2018; Pottackal et al., 2020; Pottackal, Singer, et al., 2021; Pottackal, Walsh, et al., 2021), light pulses that activated ReaChR expressed in B/K WACs evoked inhibitory postsynaptic currents (IPSCs) in RGCs (see Methods). IPSCs in OFF Delta RGCs were substantial (133 ± 46 pA; n = 11) and blocked largely by the GABA-A receptor antagonist gabazine (50 µM SR95531) (**Figure 5C**); addition of the glycine receptor antagonist strychnine (1 µM) eliminated the IPSCs completely (**Figure 5C**).

Additionally, the evoked IPSCs increased monotonically with the intensity of light stimulation and ReaChR activation (**Figure 5D**). Similar results were observed in recordings from ON Alpha RGCs: IPSCs (231 ± 32 pA; n = 19) were blocked largely by gabazine and completely blocked by addition of strychnine (**Figure 5E-H**). By comparing the ratio of early (20-120 ms) and late phases (150 – 250 ms) of the optogenetic response, we found that OFF Delta RGC responses were more transient (i.e., showed a higher early:late response ratio), whereas ON Alpha RGC responses were more sustained (**Figure 5I**).

In comparison to the IPSCs recorded in OFF Delta and ON Alpha RGCs, much smaller responses were observed in the OFF-Transient Alpha cell (28 ± 11 pA; n = 8), which stratifies between the ChAT bands and overlaps B/K WAC processes more sparsely (**Figure 5-figure supplement 1A-C**), and in DS-RGCs (18 ± 8 pA; n = 4) that co-stratify with the ChAT bands (**Figure 5-figure supplement 1D**). Thus, B/K WACs provide particularly robust synaptic input to OFF Delta and ON Alpha RGCs.

### B/K WACs make synapses with non-orientation tuned bipolar cells in the OFF layer and implement sign-inverted feedforward inhibition (SIFI)

To probe possible connections between B/K WAC dendrites and bipolar cells, we examined connectivity at the ultrastructural level. We started with an existing dataset, SBEM (scanning block-face electron microscopy) volume k0725 (Ding et al., 2016), in which all presumed inhibitory (i.e., non-ribbon) synapses onto an OFF Delta RGC had been annotated and some presynaptic amacrine cells, including WACs, had been traced (Grimes et al., 2022) (**Figure 6A, B**). The OFF Delta RGC in k0725 received 1298 inhibitory synapses, with ∼23% of these (278) coming from the narrow-field, glycinergic AII amacrine cell (Demb & Singer, 2012). Of the remaining synapses, 150 amacrine cells were reconstructed at least partially (Grimes et al., 2022), and these provided a total of 300 (out of 1020 non-AII) synapses to the OFF Delta RGC. This group comprised three classes: bistratified amacrine cells with dendrites at the borders of the INL and GCL (sublaminae S1 and S5) (Grimes et al., 2022); amacrine cells with asymmetric dendritic trees (Grimes et al., 2022); and single WAC neurites that ran unbranched through S1 of the IPL, in the same plane as the dendrites of the OFF Delta RGC (**Figure 6-figure supplement 1Bii**). The WAC neurites comprised 206 of the 300 synapses noted above, although the sampling was biased because we focused on skeletonizing processes that appeared to come from WACs.

**Figure 6:**
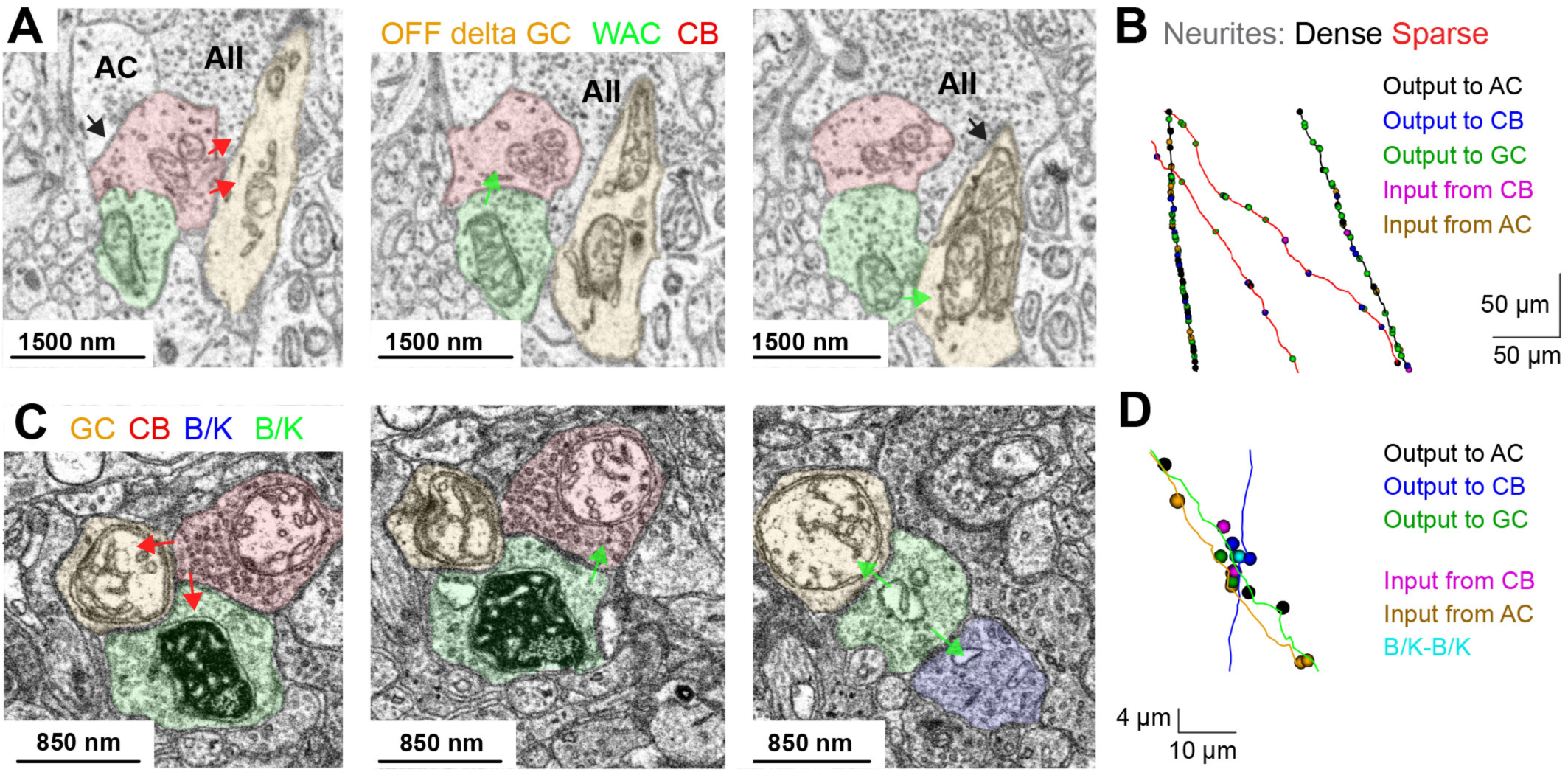
Connectomic analysis uncovers synaptic connections between B/K WACs and bipolar cells. **A.** SBEM sections from dataset k0725 show connectivity between a WAC (green), an OFF cone bipolar cell terminal (CB; red), and an OFF Delta RGC (yellow). Also shown are an unidentified amacrine cell (AC) and an AII amacrine cell (AC). Cell-type identification was determined from reconstructions in a previous study (Grimes et al., 2022). The WAC makes synapses (green arrows, middle and right panels) onto both an OFF bipolar cell terminal and the RGC dendrite that is postsynaptic to the OFF bipolar cell terminal, the signature of sign-inverted feedforward inhibition (SIFI). Additional BC synapses (red arrows) and non-WAC AC synapses (black arrows) are indicated. **B.** Sample WAC neurites (top-down view) with SIFI synapses. Contacts are color-coded per the text key. Some WACs (black lines) exhibit dense synaptic connectivity, whereas others (red lines) have sparse connectivity. **C.** SBEM sections from B/K dAPEX2 retina showing DAB^+^ processes in the OFF layer adjacent to the INL (sublamina S1). Left, OFF cone bipolar (CB, red) ribbon-type synapse (red arrows) to RGC (orange) and to DAB^+^ WAC (green). Center, DAB+ WAC inhibitory feedback synapse to CB (green arrow). Right, inhibitory synapses (green arrows) from DAB^+^ WAC to RGC and to another DAB^+^ WAC. **D.** Sample of reconstructed B/K WAC neurites (top-down view) from the B/K dAPEX2 retina with synapses color-coded per text key shows connections between two B/K WACs (green and blue dendrites) and an RGC (orange).

We quantified properties of the WAC neurites that made synapses with OFF Delta RGCs by evaluating synapse density and organization of nearby output synapses from straight, unbranched neurites. There were apparently two groups of WAC neurites with either sparse or dense total number of synapses (i.e., combined inputs and outputs) (**Figure 6-figure supplement 1Bii**). At least some of these straight neurites might belong to B/K WACs and we therefore sought to address this hypothesis to gain information about the synaptic connectivity of these cells at the ultrastructural level.

Additionally, we found that ∼25% of the inhibitory synapses to the OFF Delta RGC arose from an amacrine cell element that also provided an inhibitory synapse to a presynaptic type 2 cone bipolar cell that itself was presynaptic to the OFF delta RGC; thus, there appeared to be simultaneous pre- and postsynaptic inhibition arising at a substantial number of sites on the OFF Delta RGC dendrite. We term this synaptic motif **SIFI**, for **S**ign-**I**nverting **F**eedforward **I**nhibition (**Figs. 6A, 7C**). SIFI comprises inhibition of the RGC (WAC◊RGC) and a presynaptic excitatory (bipolar) cell (WAC◊BC◊RGC); aside from the sign reversals, this motif resembles feedforward inhibition, which comprises bipolar cell excitation of the RGC (BC◊RGC) and a presynaptic inhibitory (amacrine) cell (BC◊AC◊RGC) (**Figure 7C**).

**Figure 7.**
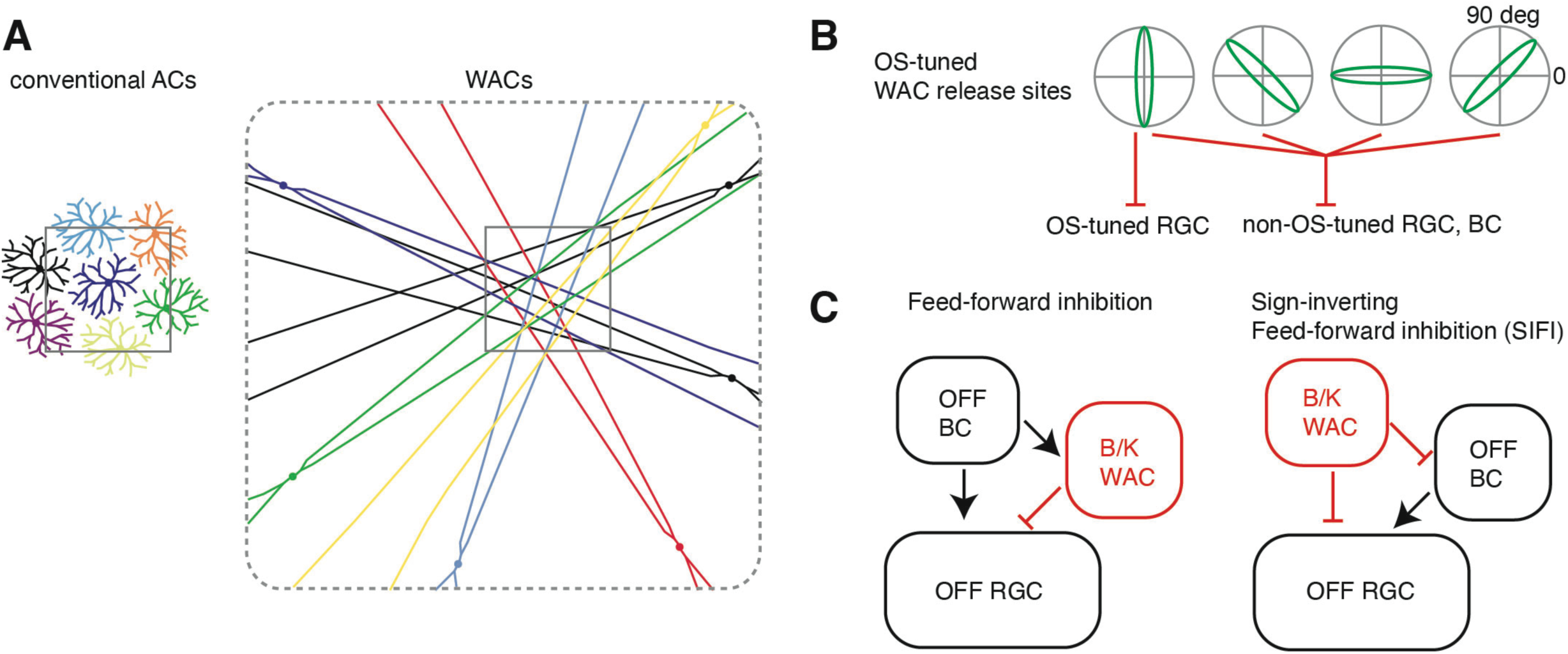
Proposed circuit function of B/K WACs. **A.** A mosaic of amacrine cells can provide coverage to a region of retina (square) with either tiled processes of seven narrow-field cells or straight dendrites projecting from the same number of WACs. **B.** Either selective or non-selective integration of OS-tuned WAC dendrites can result in either OS-tuned or non-OS-tuned postsynaptic neurons. **C.** Schematic diagrams for feed-forward inhibition and sign-inverted feed-forward inhibition (SIFI).

SIFI is found also in AII synaptic output (140/278; ∼50% of synapses), which is expected given that AIIs provide inhibitory input both to OFF Delta RGCs and to type 2 OFF cone bipolar cells presynaptic to these RGCs (Graydon et al., 2018; Tsukamoto & Omi, 2017). Notably, synapses from other amacrine types also exhibit SIFI (205/1020; ∼20%) with the OFF Delta RGC. By contrast, the SIFI motif was observed infrequently at a reconstructed OFF Alpha RGC, where only ∼8% of AII amacrine cell synapses (20/248) and ∼4% of non-AII amacrine cells (36/916) exhibited SIFI (data not shown). Thus, SIFI seems to be implemented preferentially at synapses onto the OFF Delta RGC as compared to the OFF Alpha RGC and includes synapses made by putative B/K WACs.

Because amacrine cell types cannot be identified from the SBEM volume based on morphologies alone, we performed ultrastructural analysis of a new, small volume in which B/K WACs expressed a genetically-encoded marker for use with SBEM. We targeted the enhanced peroxidase dimeric APEX2 (Joesch et al., 2016; Zhang et al., 2019) to the mitochondrial matrix of B/K WACs by dual Cre/Flp-mediated recombination and visualized genetically-identified neurites by the DAB reaction (**Figure 6C**).

Assessment of dAPEX2 expression revealed DAB+ mitochondria in varicosities that are presynaptic to presumed RGC dendrites (i.e., processes that lack synaptic vesicles) and OFF cone bipolar cell terminals in S1 (i.e., axon terminals that contained ribbon synapses and vesicles) (**Figure 6C**), although the exact cell types could not be determined in this small volume. Furthermore, some of the B/K WAC processes exhibited SIFI-connectivity (**Figure 6C, D**), matching the pattern of straight WAC neurites reconstructed from the k0725 SBEM volume. The labeled B/K WAC dendrites exhibited a relatively high density of synapses, matching the subset of WAC neurites with high synapse density reconstructed from k0725 (**Figure 6-figure supplement 1C, D**) and include outputs to cone bipolar cells (**Figure 6C, D and Figure 6-figure supplement 1Bi**). The combined analysis of optogenetic and connectomic data suggests that B/K WACs, which carry orientation-tuned signals (**Figure 4**), make output synapses to postsynaptic cells that are not themselves orientation selective.

## DISCUSSION

We developed an approach for systematic study of an enigmatic group of retinal interneurons: non-spiking wide-field amacrine cells (WACs) with straight, unbranched dendrites that are either ON or OFF cells, consistent with their stratification level in the IPL (**Figs. 1-3**). Ca^2+^ imaging showed that B/K WAC dendrites exhibited functional orientation tuning, consistent with their physical orientation (**Figure 4**). The mechanism for this tuning depends on an extended receptive field along the dendrite that is spatially isolated relative to the soma (Lei et al., 2024; Ramakrishna et al., 2026). Optogenetic studies showed that B/K WACs made output synapses onto non-orientation tuned RGC types, driving strong GABAergic inhibitory currents in ON Alpha and OFF Delta RGCs (**Figure 5**). Connectomic analyses showed that B/K WACs make synapses with bipolar cell axon terminals, which likewise lack orientation tuning (**Figure 6**) (Matsumoto et al., 2021a). The nature of encoding visual images with orientation-tuned amacrine cell dendrites suggests a novel organization for WACs. Rather than encoding local subregions via a tiled mosaic of narrow-field cells, WACs apparently overlap substantially with their neighbors and cover orientation space through the collective processing across cells with different dendrite orientations (**Figure 7A**). Networks of dendrites that are individually orientation-tuned could deliver a net tuned or non-tuned inhibition to postsynaptic neurons, depending on the pattern of convergence (**Figure 7B**). The connections shown here suggest that orientation-tuned inhibition plays an unexpected role in controlling the responses of non-orientation-tuned postsynaptic neurons in the retina (**Figs. 5, 6**).

### WACs form a diverse group of interneuron types in mammalian retina

B/K WACs share some properties with previously described cells visualized with methods for sparse labeling. Two studies identified a total of six WAC types that lacked separate dendritic and axonal compartments and can be compared to the B/K WACs: Cluster 1 and bifid cells (Badea & Nathans, 2004); and WA1, WA2-2, WA3-2 and WA4-3 cells (Lin & Masland, 2006). In comparing the soma location and dendritic tree properties, only bifid cells have properties consistent with those of B/K WACs: relatively asymmetric dendritic trees and mainly in the OFF layer with somas in the INL (Badea & Nathans, 2004). Collectively, these results suggest a diversity of WAC types in the mouse retina that extends beyond the B/K WACs described here.

Additional evidence for WAC diversity comes from both anatomical and physiological studies in mouse and other species. For example, SBEM reconstruction of mouse retina identified two groups of straight, unbranched dendrites that made synapses with one of two types of ON bipolar cell terminals (either Type 7 or Type 5A) (Hanson et al., 2023). Further, calcium signals in Type 5A bipolar cells showed orientation tuning, particularly for vertical orientations (Hanson et al., 2023). The working model for this circuit presumes that straight WAC dendrites produce orientation-tuned signals and convey this information to the Type 5A bipolar cell terminals via gap junctions; and these orientation-tuned signals were apparently further conveyed to postsynaptic DS-RGCs via glutamate synapses. The presumed orientation-tuned signals in WAC dendrites are supported by the direct evidence of OS tuning shown here (**Figure 4**). However, the B/K WACs apparently comprise different cell types than those connecting Type 5A bipolar cells, which would presumably stratify proximal to the ON ChAT band and between the ChAT bands (Hanson et al., 2023).

WACs with similar morphology to B/K cells were described as ‘wiry’ in the primate retina (human and monkey) and stratify in at least three layers: ON, OFF and near the ON/OFF boundary (Kolb et al., 1992; Manookin et al., 2015; Mariani, 1990). Similar WACs were identified in rabbit retina as well as in other species (MacNeil et al., 1999). The collected findings suggest that WACs with long, straight dendrites exist across species and across multiple layers of the retina within a species. To the extent that WACs with long, straight dendrites commonly convey orientation-tuned GABA release, this property must be fundamental across mammalian retina. Notably, OS signals are also conveyed by at least one type of spiking, axon-bearing WAC in rabbit (Murphy-Baum & Taylor, 2015), and they are also present within a narrow-field non-spiking glycinergic amacrine cell in mouse (Mani et al., 2023).

### Synaptic properties of B/K WACS

B/K WACs appear to be GABAergic based on the optogenetic response in postsynaptic RGCs, which was primarily blocked by a GABA-A receptor antagonist. But, some B/K WACs may have mixed GABA/glycine release (Lu et al., 2008; Sawant et al., 2021) (**Figure 1B**), given that residual IPSCs were blocked by the glycine receptor antagonist strychnine. This was not the case in studies of other amacrine cell types performed in other Cre lines (VIP, CRH, ChAT nNOS); in these lines, presumed GABAergic synapses, activated by stimulation of ChR2, were blocked completely by a GABA-A receptor antagonist (Park et al., 2015; Park et al., 2020; Park et al., 2018; Yonehara et al., 2011). An alternate explanation for strychnine-sensitive responses evoked by the optogenetic stimulation of B/K cells is polysynaptic activity, e.g., B/K WACs could disinhibit glycinergic cells or could excite glycinergic cells via gap junctions (Jo et al., 2023). Further studies would be required to investigate these possibilities.

The reconstruction of B/K dendrites showed a prominent pattern of SIFI, which would presumably produce a powerful form of inhibition that hyperpolarizes the RGC dendrite while simultaneously suppressing glutamate release from the presynaptic bipolar cell terminal. The SIFI synaptic motif of B/K WACs resembles the powerful glycinergic inhibition mediated by the AII amacrine cell (Demb & Singer, 2012; Ke et al., 2014), and in both cases it seems to be a specific property of input to the OFF Delta RGC but not to the OFF Alpha RGC.

### Possible roles of B/K WACs in retinal information processing

Amacrine cells with strong feature tuning to direction, motion or color are presumed to generate a similar degree of tuning in postsynaptic RGCs (Chen & Li, 2012; Sher & DeVries, 2012). For example, starburst amacrine cell pre-synapses activated by leftward motion would shape the responses of postsynaptic DS RGCs that report rightward motion (Briggman et al., 2011; Wei, 2018). Similar circuits mediate OS tuning in some RGC types in both mouse and zebrafish, where an RGC tuned to one orientation (e.g., horizontal) would receive inhibition tuned to the orthogonal orientation (e.g., vertical) (Antinucci et al., 2016; Nath & Schwartz, 2016; Venkataramani & Taylor, 2016). Electrical coupling to OS-tuned WAC dendrites also can drive OS tuning in postsynaptic bipolar cells and RGCs (Hanson et al., 2023; Nath & Schwartz, 2017).

We have shown here an additional pattern whereby OS-tuned amacrine cells could influence RGC and OFF bipolar cells that are not themselves OS-tuned. Presumably, the postsynaptic RGC or bipolar cell collects inhibitory synapses from B/K WACs of multiple orientations, preventing a strong net OS tuning in the population of connected dendrites and ensuring a lack of OS-tuning in the postsynaptic response (**Figure 7A, B**). There may be an advantage to encoding visual images with OS-tuned interneurons (**Figure 7A**), for example, since oriented features form core components of natural images (Olshausen & Field, 1996). The oriented dendrites of a B/K WAC could also promote the correlation of signaling across distant points on the retina (McIlwain, 1964; Neuenschwander & Singer, 1996).

There are other cases where tuned neurotransmitter release from presynaptic sites is canceled in the postsynaptic neuron. For example, acetylcholine released from starburst amacrine cells onto RGCs lacks strong DS tuning, because cholinergic transmission is paracrine and converges across multiple angles of direction tuning (Lee et al., 2010; Pottackal et al., 2020; Sethuramanujam et al., 2021). A similar model explains how OS-tuned glycinergic release from narrow-field VGluT3 amacrine cells could produce non-OS-tuned glycinergic inhibition of ON DS-RGCs (Mani et al., 2023).

### Limitations of the study and next steps in understanding WAC function

Our genetic approach to labeling B/K WACs has obvious advantages for targeting cells systematically. Still, though, it remains to be understood how many cell types and what fraction of an individual type is identified in any given mouse line. For example, do the B/K WACs that stratify between the ChAT bands represent a rare and sparse cell type, or are we observing sparse labeling of a dense cell type? Our inquiries therefore would benefit from additional functional tests that distinguish B/K WAC types based on Ca^2+^ imaging or by molecular markers that label soma mosaics.

In the future, we also seek methods for inactivating B/K WAC types. Inactivation studies could evaluate potential roles for B/K WACs in regulating overall excitability of RGCs and shaping OS tuning in specific RGC types. Indeed, it will be important to determine the complete set of B/K WAC postsynaptic targets and determine whether this includes OS-tuned RGCs as well as OS-tuned amacrine cells or bipolar cell release sites (Hanson et al., 2023; Matsumoto et al., 2021b; Matsumoto et al., 2025). Further, there is a possible role that WACs could play in development of other retinal cell types via temporary connections during postnatal development (Gamlin et al., 2020). By controlling B/K WAC activity more specifically, one could also learn how these cells shape responses that influence activity in the brain and behavior (Cruz-Martin et al., 2014; Hillier et al., 2017; Schroder et al., 2020).

## MATERIALS AND METHODS

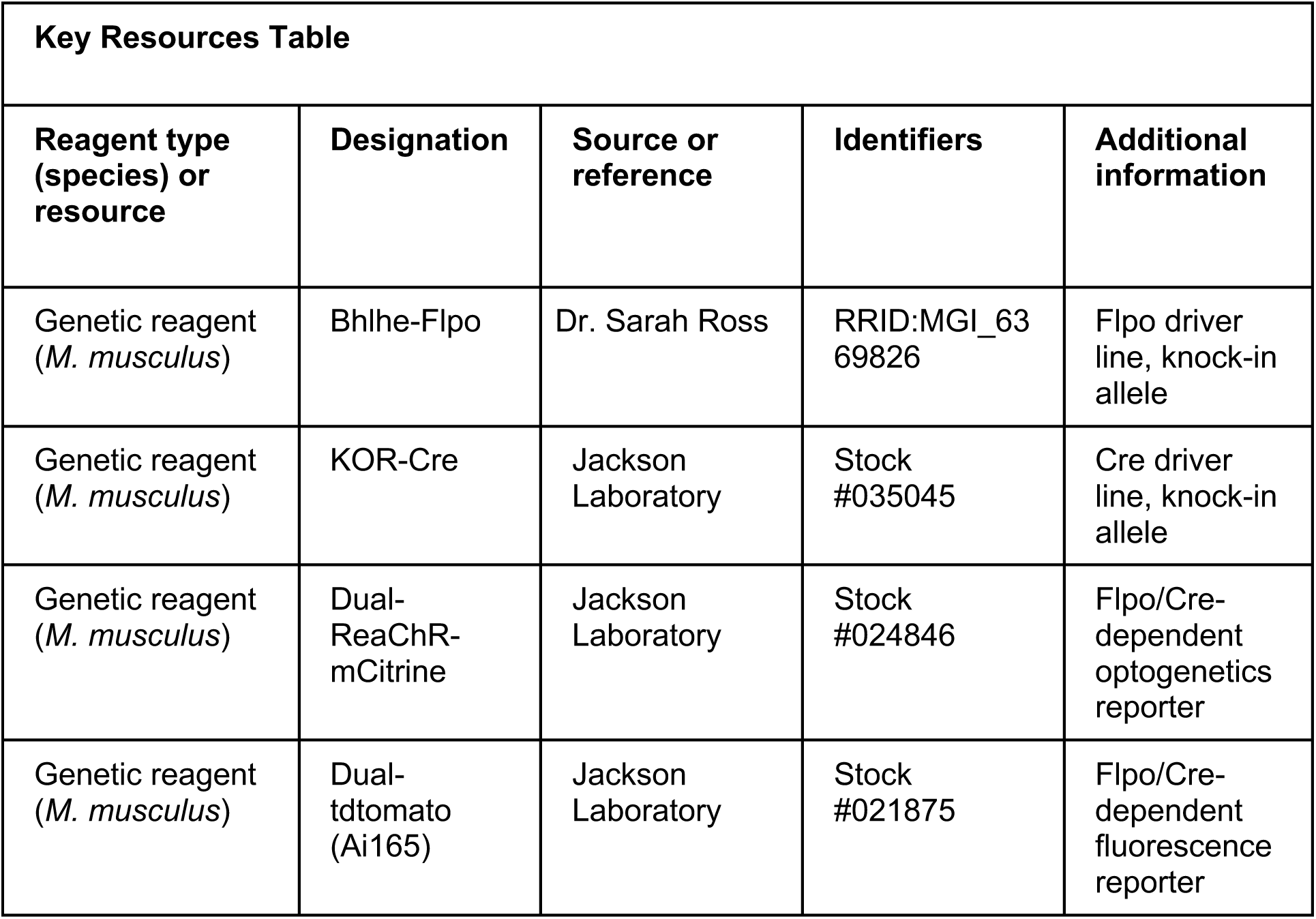
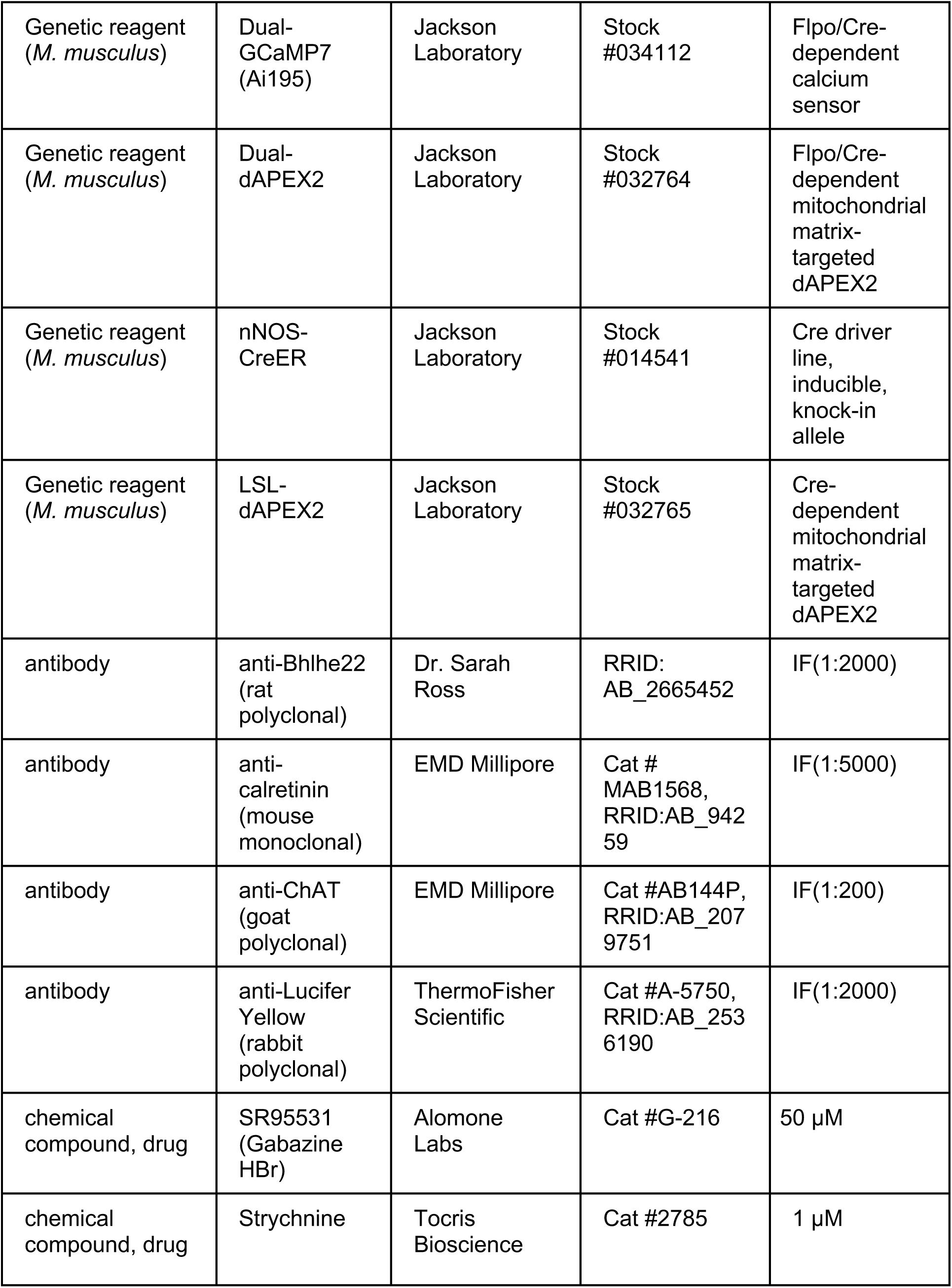
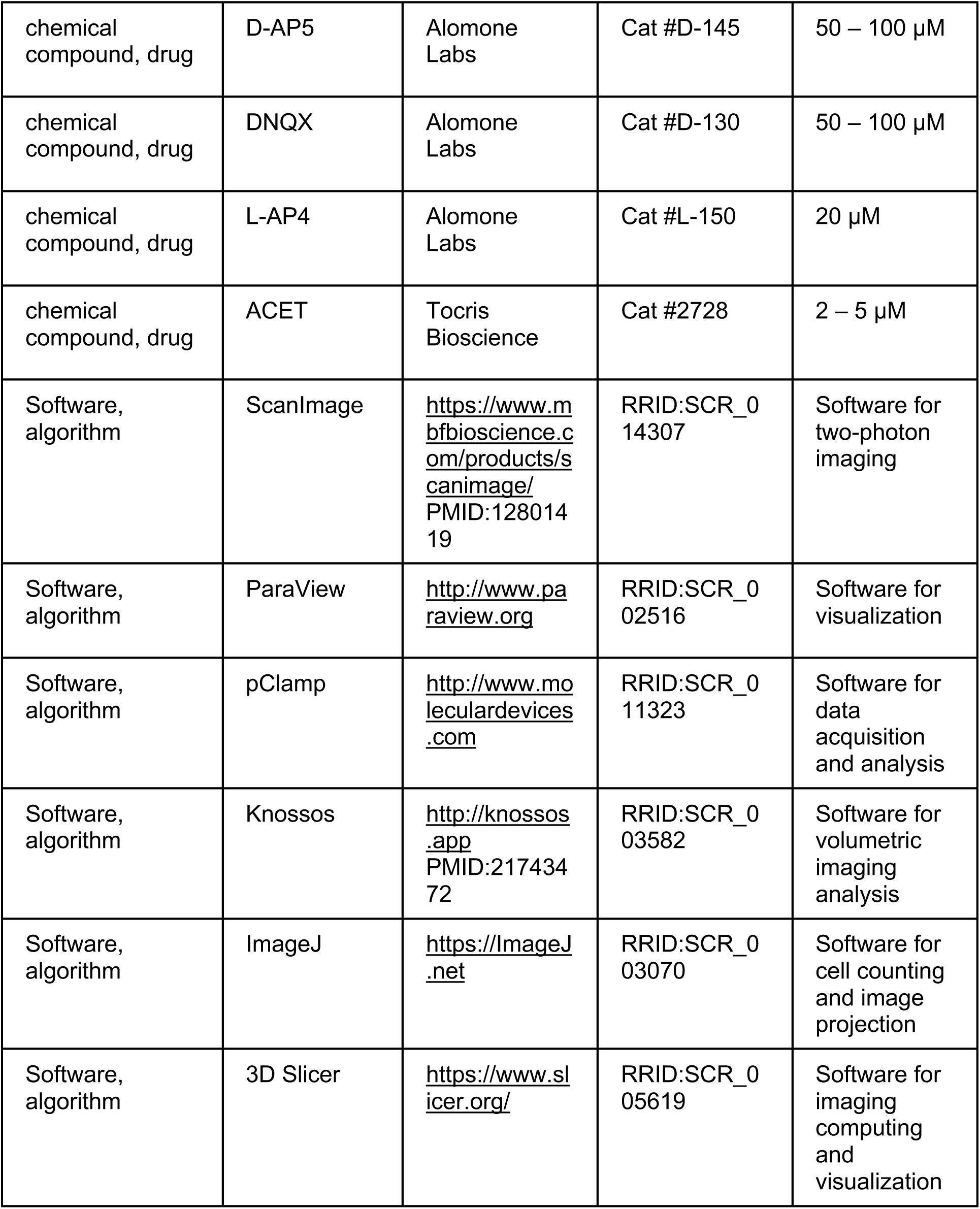

### Experimental model and subject details

All animal procedures were approved by the Institutional Animal Care and Use Committees at Yale University, University of Maryland or University of Pittsburgh and were in compliance with National Institutes of Health guidelines. For *Bbhlhe22*-Flpo knock-in/knock-out mice, Flpo recombinase was targeted to the endogenous *Bhlhe22* (*Bhlhb5*) locus, where it replaced the *Bhlhe22* gene (Cai, Kardon, et al., 2016). For KOR-Cre knock-in/knock-out mice, Cre recombinase was targeted to the endogenous KOR locus (*Oprk1*) (Cai, Huang, et al., 2016) (Jackson Laboratory #035045). Dual Cre/Flpo-dependent reporter mice included the following: dual-dependent tdTomato line (Ai65; Jackson Laboratory #021875), dual-dependent ReaChR-mCitrine fusion protein line (Rosa26 CAG-LoxStopLox-FrtStopFrt-ReaChR-mCitrine; Jackson Laboratory #024846), dual-dependent GCaMP7s line (Ai195; Jackson Laboratory #034112); and dual-dependent mitochondrial matrix-targeted dimeric APEX2 line (ROSA26^DR-Matrix-dAPEX2^; Jackson Laboratory #032764).

Mice were bred to have appropriate combinations of Cre, Flpo and reporter alleles. In the cases of KOR-Cre and *Bhlhe22*-Flpo alleles, all mice studied were heterozygous for each transgenic allele and therefore had one intact copy of each gene. Pilot studies that crossed either KOR-Cre with a Cre-dependent reporter line (Ai32) or *Bhlhe22*-Flpo mice with a Flp-dependent reporter line (Cai, Kardon, et al., 2016) suggested that both lines contained WACs with straight, unbranched dendrites, which motivated the intersectional approach described above. All mice were maintained on a 12/12-hour light/dark cycle and were studied at 1.5-6 months of age. We did not expect or detect differences between male and female mice, and results have been combined across sex.

### Electrophysiology and Cell Filling

Prior to retinal dissection, a mouse was dark-adapted for ∼1 h. After death, both eyes were enucleated and maintained in Ames medium (MilliporeSigma) supplemented with 22.6 mM NaHCO_3_ (MilliporeSigma) and suffused with 95% O_2_/5% CO_2_. The dissection was performed under infrared illumination using stereomicroscope-mounted night-vision goggles (OWL Night Vision Scopes, third generation; B. E. Meyers). The retina was removed from the eyecup, the vitreous humor was removed, and one or more relaxing cuts were made in order to flatten the retina. The retina was mounted on a filter membrane (HAWP01300, MilliporeSigma) with five 0.5-mm diameter holes punched out and maintained in a dissection dish at room temperature for up to several hours. The orientation of the retina was not tracked precisely in all cases, and hence we did not track cell orientation relative to the body axis. For recording, the retina was placed into a custom recording chamber and secured with a tissue harp. The recording chamber was perfused with Ames medium flowing at 4-6 mL min^-1^ and maintained at 31 – 33 deg C.

Whole-cell patch-clamp recordings were obtained using patch pipettes pulled from borosilicate glass capillaries (1B120F-4, World Precision Instruments). Pipette tip resistances were ∼4-10 MΩ for amacrine cell recordings in the ganglion cell layer; ∼7-10 MΩ for amacrine cell recordings in the inner nuclear layer; and 2-10 MΩ for ganglion cell recordings. Pipettes were filled with internal solution containing (in mM): 120 K-methanesulfonate, 10 HEPES, 0.1 EGTA, 5 NaCl, 4 ATP-Mg, 0.4 GTP-Na_2_, and 10 phosphocreatine-Tris_2_ (pH 7.3, 280 mOsm) for current-clamp recordings and 120 Cs-methanesulfonate, 5 TEA-Cl, 10 HEPES, 10 BAPTA, 3 NaCl, 2 QX-314-Cl, 4 ATP-Mg, 0.4 GTP-Na2, and 10 phosphocreatine-Tris2 (pH 7.3, 280 mOsm) for voltage-clamp recordings. The internal solution also contained 0.05% (w/v) Lucifer Yellow to label cells for subsequent immunohistochemical experiments. The components of the internal solution were obtained from MilliporeSigma. Membrane potential or current was amplified (MultiClamp 700B, Molecular Devices), digitized at 10 kHz (Digidata 1440A, Molecular Devices) and recorded (pClamp 10.0, Molecular Devices). Series resistance was ∼20 – 40 MΩ for amacrine cell recordings in the GCL; ∼20 – 100 MΩ for amacrine cell recordings in the INL; and ∼8-25 MΩ for ganglion cell recordings. Series resistance was typically compensated by ∼50%. Recordings were corrected for a -9 mV liquid junction potential.

### Two-photon imaging

Two-photon fluorescent measurements were made using a custom-built microscope (Olympus BX-51) equipped with an Olympus 60×, 0.9 NA, LUMPlanFl/IR water immersion objective (Olympus), two fluorescence detection channels for GCaMP7s (HQ 535/50, Chroma) and SR101 (DM 565, 650/SP, Semrock), and an ultrafast pulsed laser (Chameleon Ultra II; Coherent) tuned to 910 nm. ScanImage software version 3.8.1 was used for image acquisition. Time-lapsed images (128 × 128 pixels; 26.8 × 26.8 μm frame) at each IPL depth were acquired at 7.8 frames per second with imaging laser power less than 30 mW. For volumetric imaging in the IPL, 12 time-lapsed z-stacks with 4-μm spacing were captured to cover the B/K WAC dendrites in the IPL. The z plane on the distal side (i.e., towards the INL) of the dendrites in the ON layer was defined as plane 0 for aligning across z stacks.

### Visual stimulation

Visual stimuli were generated using custom software written in either Stim-Demo (C language) (Borghuis et al., 2014; Borghuis et al., 2013) or in MATLAB using the Psychophysics Toolbox (Brainard, 1997) and were displayed by a DLP projector (HP Notebook Projection Companion) with 60-Hz refresh rate, modified to project UV light (peak wavelength at 395 nm; Borghuis et al., 2013). This wavelength about equally activates cone photoreceptors along the retina’s dorsal/ventral axis, which have varying ratios of green- and UV-sensitive opsins (Wang et al., 2011). The image of the DLP projector was focused on the retina through the microscope condenser.

Stimuli were presented within a 4 x 3 mm area on the retina and included contrast-reversing spots of variable diameter and drifting sine-wave gratings, both at 100% Michelson contrast relative to the background mean luminance that evoked ∼10^4^ photoisomerizations cone^-1^ s^-1^. For the drifting grating stimuli used for calcium imaging, a full-field background of mean luminance was presented for 5 s to let the retina adapt to the combination of laser and mean luminance, and subsequently sinusoidal gratings moving in eight directions were presented sequentially. Each grating stimulus was presented for 2 s with a 5 s interval between stimuli. The grating was presented in a patch of 880 µm (29.3 degrees) in diameter and had a 2-Hz temporal frequency and 0.139 cycles/degree (4.54 cycles/mm) spatial frequency.

Spot tuning experiments were performed with spots of diameter 50 – 1675 µm. Because slightly different spot sizes were presented across experiments, responses were binned based on spot diameters centered around 100 µm (50 – 275 µm), 450 µm (276 – 625 µm), 800 µm (626 – 975 µm), 1150 µm (976 – 1325 µm) and 1500 µm (1326 – 1675 µm) before averaging across cells.

### Imaging data analysis

Calcium imaging data were analyzed using custom software written in MATLAB. Data were analyzed within regions of interest (ROIs). To select ROIs, a Gaussian-blurred mean image was first subtracted from each frame for background fluorescence subtraction. Then, a local correlation image was calculated. Each pixel in the local correlation image was the mean of the correlation between the center pixel and the neighboring pixels in the surrounding 3x3 matrix. Rectangular ROIs were manually selected from non-overlapping dendritic segments on the correlated image, with one ROI per isolated dendritic segment. The ROIs were drawn as rectangles with the long side of the rectangle parallel to the dendrite.

Next, the raw fluorescence and physical orientation *θ_p_*_ℎ*y*_ of each ROI were extracted based on the binary ROI mask. For the calcium response to the drifting gratings, *R*(*θ_k_*) is calculated as the rectified mean of the baseline-subtracted fluorescence evoked by the grating moving in the direction *θ_k_* for each ROI or image pixel. The baseline was defined as the average fluorescence 2.5 - 5 seconds after laser onset and just before stimulus onset.

An orientation selective index (OSI) was defined as follows:

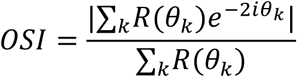

which is equivalent to 1 − circular variance in circular statistics. Note that the factor of 2 in the OSI equation makes angles separated by 180° equivalent, so the measure quantifies orientation rather than direction.

A direction selective index (DSI) was defined as follows:

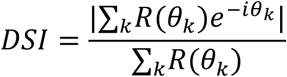

The preferred orientation was calculated as follows:

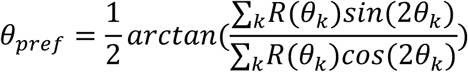

The circular Pearson correlation (*r_circ_*) between *θ_p_*_ℎ*y*_ and *θ_pref_* in orientation space was defined as follows:

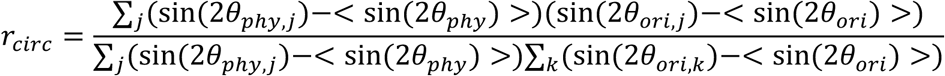

To demix spatially overlapping dendrites and visualize spatial components of apparent dendritic structures that were otherwise indistinguishable in the correlation image, we utilized non-negative matrix decomposition to decompose 3d video *Y*(*t*) into products of spatial footprint *A_i_* and corresponding temporal fluorescence traces *C_i_* and background noise *B*(*t*):

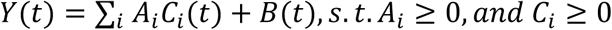

using the alternative least square algorithm (Berry et al., 2007).

### Optogenetics

For optogenetic experiments, conventional photoreceptor-mediated input was pharmacologically blocked via bath application of the following drug cocktail (in µM): 50-100 D-AP5 (Alomone Labs), 50-100 DNQX (Alomone Labs), 20 L-AP4 (Alomone Labs) and 2-5 ACET (Tocris) (Park et al., 2015; Park et al., 2020; Park et al., 2018; Pottackal et al., 2020; Pottackal, Singer, et al., 2021; Pottackal, Walsh, et al., 2021). ReaChR-mediated responses were evoked by an LED (λ peak = 450 or 470 nm; maximum intensity ∼5 x 10^17^ photons s^-1^ cm^-2^) focused through the condenser onto a square (220 µm side) as described (Park et al., 2015; Park et al., 2020; Park et al., 2018) and presented for a 250-msec duration (average of 4 pulses, 1-s inter-pulse interval, repeated three times). The efficiency of ReaChR stimulation with blue light is within a factor of two as compared to the peak sensitivity using red light (Lin et al., 2013).

Optogenetically-evoked responses were measured as postsynaptic currents in RGCs. The response amplitude was the average current over 120 msec surrounding the peak response minus the baseline current, measured over 200 msec before light onset. In a separate analysis of response timing, amplitudes were compared in early (20 – 120 ms) and late (150 – 250 ms) time windows. Optogenetic stimulation can evoke melanopsin-mediated responses in ON Alpha RGCs, but these responses run down over time and would be very weak at the inhibitory reversal potential used in voltage-clamp experiments here (Beier et al., 2013).

### Histology

For whole-mount staining of filled cells, the retina was fixed after recording with 4% PFA for 1 h at room temperature. The retina was incubated in 6% donkey serum and 0.5% triton X-100 in PBS for 1 h at room temperature; and then incubated with 2% donkey serum and 0.5% Triton X-100 in PBS with primary antibodies for 1 day at 4° C, and with secondary antibodies for 2 h at room temperature.

For staining with the Bhlhe22 antibody, the retina was fixed in 4% paraformaldehyde in PBS for 1 hour at room temperature; incubated in blocking solution overnight at 4°C; in primary antibody solution for 5 days at 4°C; and in secondary solution for 2 days at 4°C. Retinas were washed three times in PBS for 20 minutes each at room temperature after primary and secondary antibodies were applied. For imaging, the retinas were mounted on slides in Fluoromount G (Electron Microscopy Sciences) with the retinal ganglion cell layer facing the cover slip.

For staining retinal sections, eyecups were placed in 4% paraformaldehyde in PBS for 1-2 hours at room temperature. Next, eyecups were rinsed twice in PBS and embedded in 7% low-melt agarose (Precisionary, SKU VF-AGT-VM). The agarose-embedded eyecups were Vibratome-sectioned into 100 μm-thick slices (VT1200; Leica) and blocked in PBS containing 5% donkey serum and 0.5% Triton X-100 for 1 hour at room temperature. Sections were incubated with primary antibodies overnight at 4°C, washed three times with PBS for 10 minutes, and incubated for 2 hours at room temperature with appropriate secondary antibodies. Stained slices were washed three times in PBS for 10 minutes, mounted onto slides in Fluoromount G (Electron Microscopy Sciences) and cover slipped.

Primary antibodies were used at the following concentrations: rat anti-Bhlhe22 (Bhlhb5; 1:2000) (Ross et al., 2010), goat anti-ChAT (1:200, Millipore AB144P, RRID:AB_2079751), rabbit anti-Lucifer Yellow (1:2000, ThermoFisher Scientific A-5750, RRID:AB_2536190), mouse anti-calretinin (1:5000, Millipore MAB1568). Secondary antibodies were conjugated to Alexa Fluor 488, C3 and C5 (Jackson ImmunoResearch or ThermoFisher Scientific) and diluted at 1:500.

### Confocal imaging and analysis

For both retinal sections and flat mounts, confocal images were taken with a Zeiss LSM800 laser scanning confocal microscope. For filled cells, a whole-mount image of the dendritic tree was acquired using a 10X (NA = 0.3) or 20X air objective (NA = 0.8). A high-resolution z-stack was obtained to determine the relative depth of the filled dendrites relative to the ChAT bands, using a 40X oil objective (NA = 1.4); a digital zoom of 0.8 was used to expand the imaged area. Given the NA, the excitation wavelengths and the pinhole setting (1 Airy unit), integration along the z-axis would be ∼0.5 µm; the z-axis spacing in 40X confocal z-stacks was 0.41 µm. Custom software written in MATLAB (MathWorks) was used to determine dendrite stratification relative to the ChAT bands. The program and methods used were similar to those described previously (Beaudoin et al., 2019; Manookin et al., 2008; Park et al., 2015). The depth of labeled processes is reported relative to the fluorescence peaks of the ChAT bands, in the *z* dimension, aligned to 0 (peak of inner ChAT band) and 1 (peak of outer ChAT band) in normalized units. In this coordinate system, the boundaries of the nuclear layers are approximately -1.2 (GCL) and 1.8 (INL) (Park et al., 2015; Park et al., 2018). In one experiment, we labeled three bands in retinal sections with the calretinin antibody (Figure 1C); the outer calretinin bands correspond to the ChAT bands (Haverkamp & Wassle, 2000).

For the antibody staining in retinal flat mounts and sections, the 40X oil objective was used, and image analysis and processing were performed with *ImageJ* software (Fiji: ImageJ2 Version 2.3.0/1.53q).

### Analysis of dendritic tree orientation bias

To analyze the orientation bias of a BK WAC dendritic tree, we modified the OSI analysis, described above for fluorescence measurements and previously to characterize cortical neuron responses (Liang et al., 2023; Niell & Stryker, 2008). We considered every dendrite to have an *R* value of 1, such that the morphological OSI (mOSI) equation simplifies to the following:

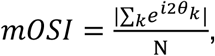

where N is the number of dendrites. To evaluate the mOSI value relative to the null hypothesis, we generated Monte Carlo null distributions of mOSI values for each relevant number of dendrites observed in the BK WAC sample (4-13 dendrites). For each number of dendrites (d_n_), we generated 10,000 samples with dendrite angles drawn independently from a uniform distribution over 0-360°. For each cell, we calculated a one-sided p-value as the fraction of simulated mOSI values greater than or equal to the observed mOSI, using the null distribution with the same number of dendrites. Under the null hypothesis, the p-value distributions would follow a uniform distribution, and we evaluated the sample p-value distribution, either for all BK WACs or from the ON BK WACs alone, relative to the uniform distribution using a Kolmogorov-Smirnov test (one-sample test).

### Connectomic analysis with Scanning Block Face Electron Microscopy

Reconstructions were performed in an existing mouse retina volume k0725 (50×210×260 μm; voxel size 13.2×13.2×26 nm) at home.webknossos.org (110629_k0725) (Ding et al., 2016); and with a new SBEM volume with labeled B/K WAC processes (61×24×40 μm; voxel size 5×5×40 nm). Manual skeletonization and annotation were performed using Knossos (Max Planck Institute for Medical Research; Heidelberg, Germany) (Helmstaedter et al., 2011). For analyses using k0725, voxel coordinates were tilt-corrected and normalized to the positions of the ON and OFF Starburst amacrine cells. Connectivity analysis was performed using custom-written Python scripts, and data were visualized using Paraview (Kitware, Inc.) (Graydon et al., 2018; Grimes et al., 2022; Park et al., 2020). For the identified OFF Delta cell, we assumed that non-ribbon synapses were inhibitory. This does not account for the vGluT3 amacrine cell, which releases glutamate but lacks ribbons (Friedrichsen et al., 2024; Lee et al., 2014); however, reconstructed vGluT3 cells could be recognized from their synapses with OFF (transient) Alpha RGCs and did not make similar synapses onto OFF Delta cells.

To prepare the new SBEM volume with labeled B/K WAC processes, a *Bhlhe22*-Flpo;KOR-Cre;dual-dAPEX2 mouse was euthanized and the retina was immersed for 30 minutes in 0.2M cacodylate buffer (pH 7.4) containing 4% paraformaldehyde, 2% glutaraldehyde, and 4 mM CaCl2. Tissue was washed with 0.2M cacodylate buffer on ice (4×20 min), immersed for 30 min in 0.2 M cacodylate buffer containing 20 mM glycine on ice, and washed with 0.1M cacodylate buffer (pH 7.4) on ice (5×10 min). To visualize APEX2 expression, tissue was immersed for 30 minutes in 0.2 M cacodylate containing 0.5 mg/ml diaminobenzidine (DAB) on ice and protected from light. H_2_O_2_ was added to a final concentration of 0.03% and tissue was incubated for 20 minutes.

Finally, tissue was washed in 0.2 M cacodylate buffer (pH 7.4) on ice (3×1 minutes, then 2×20 minutes). To prepare the tissue for SBEM, the sample was post-fixed at room temperature with 2% osmium tetroxide in 0.15 M cacodylate containing 2.5% potassium ferrocyanide, washed in three changes of distilled H_2_O, incubated at 60° C in 1% aqueous thiocarbohydrazide, washed again in distilled H_2_O, incubated in distilled H_2_O containing 1% aqueous uranyl acetate and 25% ethanol, rinsed in distilled H_2_O, incubated in lead aspartate at 60° C, and rinsed in distilled H_2_O. Then, dehydration was performed in a graded series of ethanol (35–100%) and infiltration in a propylene oxide/epoxy resin series was followed by embedding and polymerization in epoxy resin. Because of the limited volume of the newly-generated dataset, we could not identify specific RGC types.

We validated the approach above by labeling nNOS-1 amacrine cells, which are presynaptic to AII amacrine cells (Park et al., 2020). We generated a *nNOS*-CreER;LSL-dAPEX2 mouse (Jackson Laboratory #014541; Rosa 26 LSL-Matrix-dAPEX2; Jackson Laboratory #032765), administered tamoxifen (Park et al., 2020), and targeted the enhanced peroxidase dimeric APEX2 (Joesch et al., 2016; Zhang et al., 2019) to the mitochondrial matrix of nNOS+ amacrine cells by Cre-mediated recombination. We then visualized genetically-identified neurites by the DAB reaction, as described above (**Figure 6-figure supplement 2**). Assessment of dAPEX2 expression revealed DAB+ mitochondria in varicosities that are presynaptic to AII amacrine cells (**Figure 6-figure supplement 2**), confirming previously identified synapses in unlabeled tissue (Park et al., 2020).

### Transcriptomic analysis

Transcriptomic data came from a published study of single-cell RNA sequencing of amacrine cells (Yan et al., 2020). The dataset (SCP919), available from the Broad Institute Single Cell Portal (Tarhan et al., 2023), was downloaded and analyzed in custom software written in Matlab (https://singlecell.broadinstitute.org/single_cell/study/SCP919). The distributions of *Bhlhe22* and *Oprk1* transcript readings were analyzed for each of the 63 previously assigned amacrine cell groups (i.e., presumed cell types).

### Quantification and Statistical Analysis

Data are reported as mean ± SEM. P-values were calculated based on a two-tailed Student’s T-test (paired or unpaired as appropriate). No statistical methods were used to predetermine sample sizes.

## Acknowledgements

This research was supported by NIH grants R01 EY029323 (J.B.D.), R01 EY014454 (J.B.D.), R01 EY017836 (J.H.S.), P30 EY026878 (J.B.D.), T32 EY022312 (Z. Jimmy Zhou), DC-IDDRC P50HD105328 (C.C.-P., A.P.), R24 MH114785 (A.P.), F32 EY032389 (C.H.-G.), F31 EY034776 (T.E.Z.); a National Science Foundation Graduate Fellowship (J.P.) and a Gruber Science Fellowship (J.P.).

## Author contributions

Conceived the research plan: J.B.D., S.E.R., J.H.S.; Designed the experiments: S.J.P., W.L., J.P., S.E.R., J.H.S., J.B.D.; Collected and analyzed the data: S.J.P., W.L., J.P., A.O., J.M., J.P., C.H.-G., T.E.Z., C.C.-P., A.P.; Population dendritic imaging: W.L. Wrote the paper, original draft: J.B.D., W.L., J.H.S.; Edited the paper: T.E.Z., S.E.R., S.J.P.

## Resource Availability

All data, code, images and analyses have been archived and are available upon reasonable request. Further information and requests for resources and reagents should be directed to Jonathan Demb.

## Supplementary Figure Legends

**Figure 1-figure supplement 1. Transcript readings for *Oprk1* showed weak signals and were difficult to interpret.**

**A.** *Oprk1* transcript readings in previously identified groups of amacrine cells (Yan et al., 2020). Data points show the median *Oprk1* transcript reading and the inter-quartile range (IQR) of these readings within each group. Cell groups are identified as GABAergic, glycinergic, both or neither, according to the definitions in the original study, and groups that correspond to known cell types are labeled with the name of the type (e.g., AII or A17 amacrine cell) or a characteristic gene name (e.g., VIP^+^ or nNOS^+^ amacrine cells).

**B.** Same as A. for the average *Oprk1* transcript reading.

**C.** Same as A. for the percent of cells in each group with a non-zero *Oprk1* transcript reading.

**D.** The log_10_ number of cells in each group. Data for cell groups with higher numbers are based on fewer cells.

**Figure 4-video 1. B/K WAC dendrites in the OFF layer show orientation-tuned calcium signals.**

Movie shows calcium signals (GCaMP7s) in B/K WAC dendrites in the OFF layer during the presentation of gratings with various orientations and directions of motion. Movie is played at 3x original speed. Drifting grating stimuli were sinusoidal gratings moving in eight directions. Each stimulus was presented for 2 s with a 5 s interval between stimuli. The grating had a 2-Hz temporal frequency, a 0.139 cycles/degree (4.54 cycles/mm) spatial frequency and was presented in a patch of 880 µm (29.3 degrees) in diameter.

**Figure 4-video 2. B/K WAC dendrites in the ON layer show orientation-tuned calcium signals.**

Same format as Figure 4-video 1 for B/K WAC dendrites in the ON layer.

**Figure 5-figure supplement 1. B/K amacrine cells make weak connections with certain retinal ganglion cell types**

**A.** Dendritic tree of a recorded OFF Alpha RGC (also known as a Transient OFF Alpha RGC). Image shows average fluorescence in a confocal stack. Dashed square shows region in B. The RGC was filled with Lucifer Yellow (LY) during whole-cell recording, which was subsequently amplified with LY primary antibody and a red secondary antibody.

**B.** Single confocal section showing OFF Alpha RGC dendritic tree relative to B/K WACs (Cre/Flpo-dependent ReaChR-mCitrine reporter).

**C.** OFF Alpha RGCs had weaker IPSCs relative to OFF Delta and ON Alpha RGCs (Figure 5). Responses were only weakly blocked by gabazine; in one cell, there was an inward current following addition of strychnine (inset).

**D.** ON-OFF Direction-Selective RGCs showed weak IPSCs following optogenetic stimulation of B/K cells.

**Figure 6-figure supplement 1. Studying labeled B/K amacrine cells with a scanning block face electron microscopy (SBEM) data set**

**A.** Relative size and depth of the two SBEM datasets: an existing large dataset (k0725) and a smaller new data set with labeled B/K processes (B/K dApex2). The B/K dAPEX2 dataset focused on the OFF layer.

**Bi.** Sample of reconstructed B/K WAC neurites (top-down view) from the dAPEX2 retina SBEM dataset with synapses color-coded per the text key.

**Bii.** Sample WAC neurites (top-down view) from the k0725 dataset with SIFI synapses. Contacts are color-coded per the text key.

**C.** There was an apparent bias to observe higher synapse density in short segments in the B/K dAPEX2 dataset. We analyzed segments greater than 10 µm in length.

**D.** Neurites in sublamina S1 had similar densities of input and output synapses in the new data set (B/K dApex2) and the subset of straight dendrites with dense synapses in the existing data set (k0725; boxed region).

**Figure 6-figure supplement 2. dAPEX2-labeled process of an nNOS-1 amacrine cell makes a synapse with an AII amacrine cell.**

**A.** SBEM section from an nNOS-CreER;LSL-dAPEX2 retina showing a DAB^+^ process in the ON layer adjacent to rod bipolar cell terminals. The labeled process belongs to a presumed nNOS-1 amacrine cell.

**B.** Section nearby the one in A. showing a synapse from the DAB^+^ process to an AII amacrine cell, which was identified based on its morphology and its input from a rod bipolar cell.

**C.** Section nearby the one in B. showing a rod bipolar cell making a ribbon synapse with the AII amacrine cell in B.

**D.** 3D reconstruction showing the synapses illustrated in B. and C. and the labeled (DAB^+^) mitochondrion in A.

