## Supplementary material for "Molecular identification of wide-field amacrine cells in mouse retina that encode stimulus orientation": Figure 1-figure supplement 1

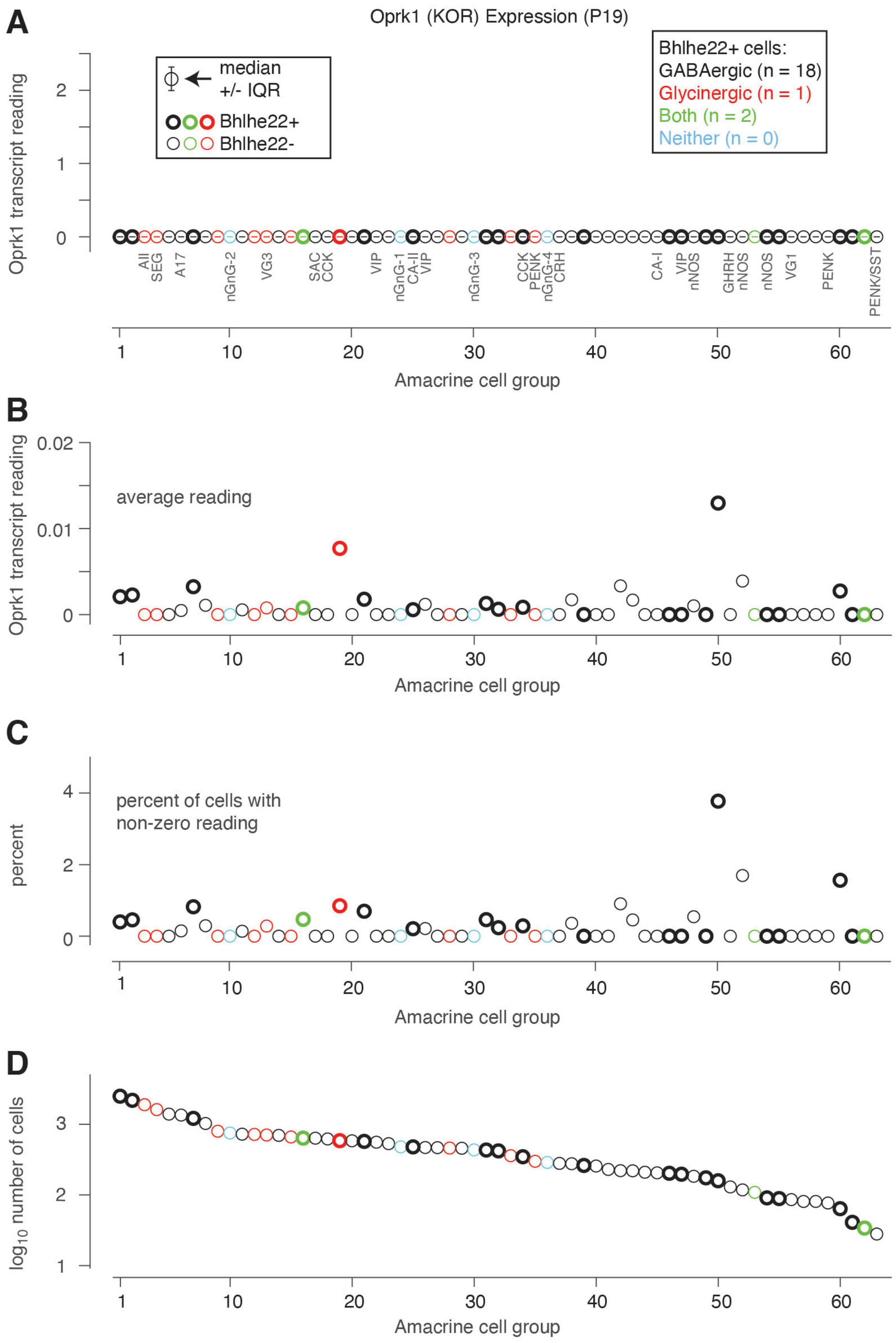


Figure 1-figure supplement 1. Transcript readings for Oprk1 showed weak signals and were difficult to interpret.

A. Oprk1 transcript readings in previously identified groups of amacrine cells (Yan et al., 2020). Data points show the median Oprk1 transcript reading and the inter-quartile range (IQR) of these readings within each group. Cell groups are identified as GABAergic, glycinergic, both or neither, according to the definitions in the original study, and groups that correspond to known cell types are labeled with the name of the type (e.g., AII or A17 amacrine cell) or a characteristic gene name (e.g., VIP+ or nNOS+ amacrine cells).

B. Same as A. for the average Oprk1 transcript reading.

C. Same as A. for the percent of cells in each group with a non-zero Oprk1 transcript reading.

D. The log10 number of cells in each group. Data for cell groups with higher numbers are based on fewer cells.
