## Supplementary material for "Molecular identification of wide-field amacrine cells in mouse retina that encode stimulus orientation": Figure 5-figure supplement 1

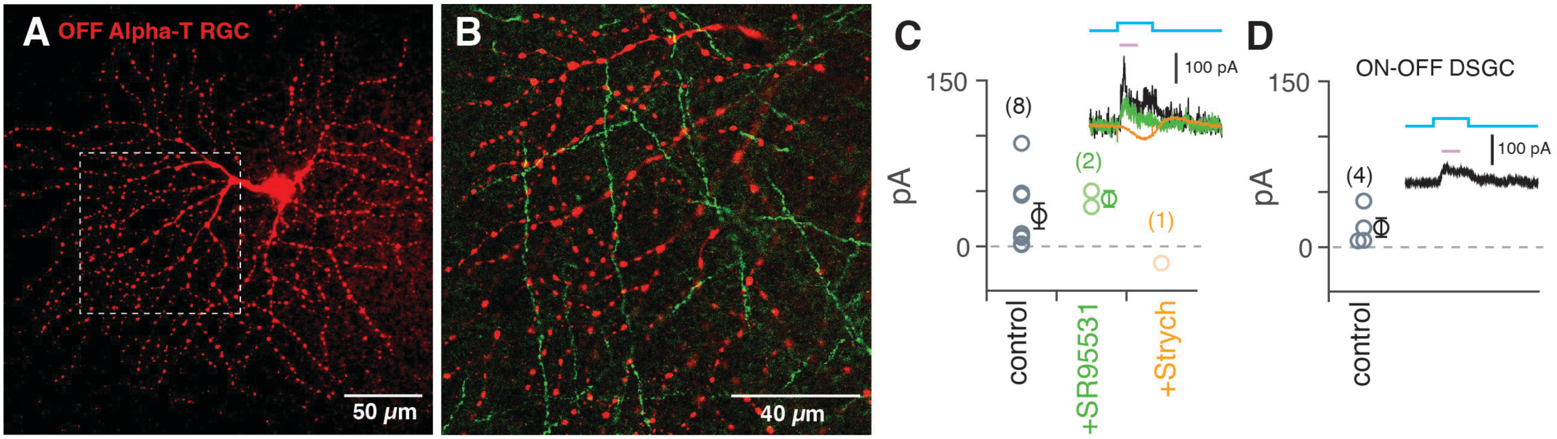


Figure 5-figure supplement 1. B/K amacrine cells make weak connections with certain retinal ganglion cell types
A. Dendritic tree of a recorded OFF Alpha RGC (also known as a Transient OFF Alpha RGC). Image shows average fluorescence in a confocal stack. Dashed square shows region in B. The RGC was filled with Lucifer Yellow (LY) during whole-cell recording, which was subsequently amplified with LY primary antibody and a red secondary antibody.
B. Single confocal section showing OFF Alpha RGC dendritic tree relative to B/K WACs (Cre/Flpo-dependent ReaChR-mCitrine reporter).
C. OFF Alpha RGCs had weaker IPSCs relative to OFF Delta and ON Alpha RGCs (Figure 5). Responses were only weakly blocked by gabazine; in one cell, there was an inward current following addition of strychnine (inset).
D. ON-OFF Direction-Selective RGCs showed weak IPSCs following optogenetic stimulation of B/K cells.
