## Supplementary material for "Molecular identification of wide-field amacrine cells in mouse retina that encode stimulus orientation": Figure 6-figure supplement 1

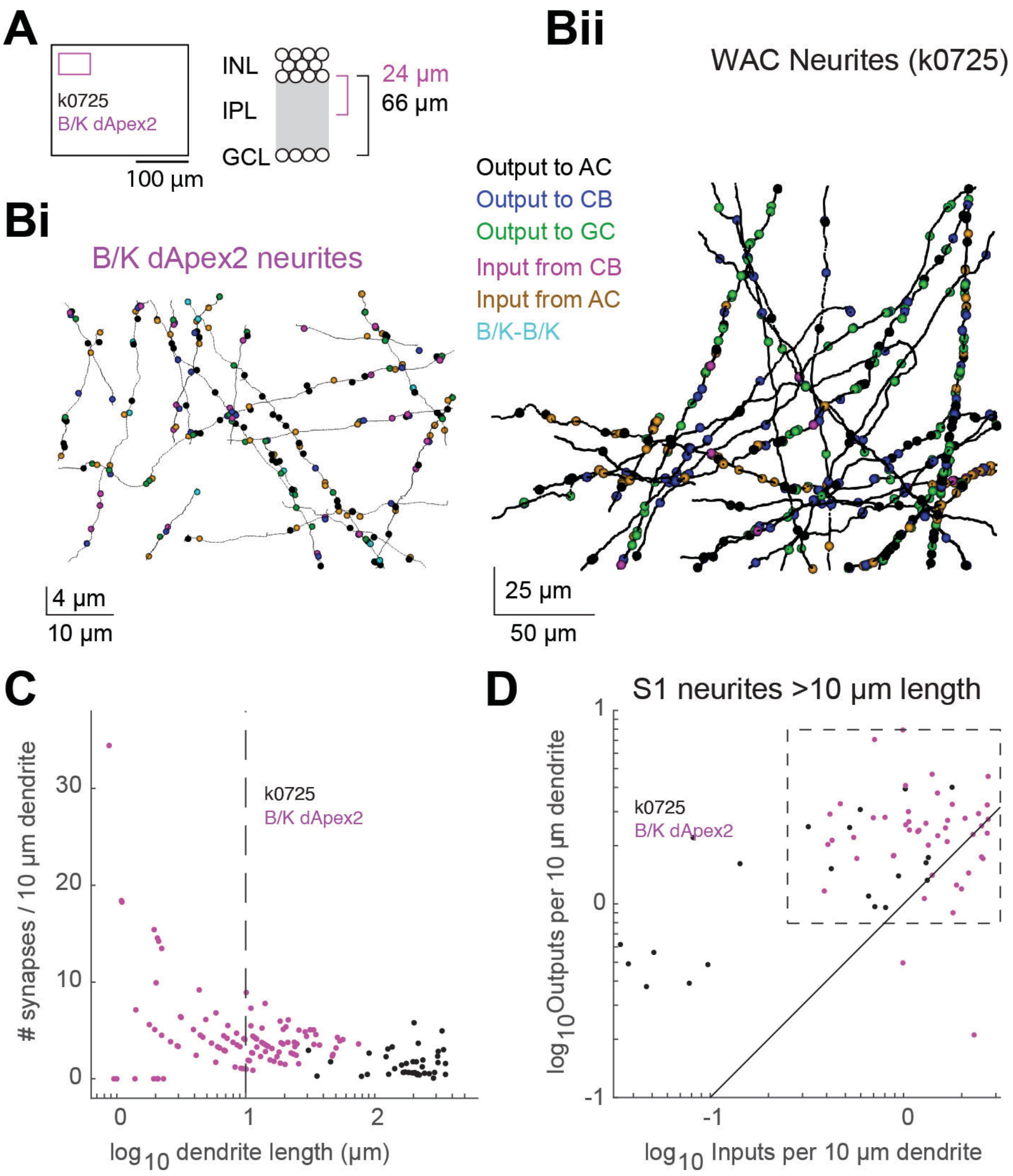


Figure 6-figure supplement 1. Studying labeled B/K amacrine cells with a scanning block face electron microscopy (SBEM) data set

A. Relative size and depth of the two SBEM datasets: an existing large dataset (k0725) and a smaller new data set with labeled B/K processes (B/K dApex2). The B/K dAPEX2 dataset focused on the OFF layer.

Bi. Sample of reconstructed B/K WAC neurites (top-down view) from the dAPEX2 retina SBEM dataset with synapses color-coded per the text key.

Bii. Sample WAC neurites (top-down view) from the k0725 dataset with SIFI synapses. Contacts are color-coded per the text key.

C. There was an apparent bias to observe higher synapse density in short segments in the B/K dAPEX2 dataset. We analyzed segments greater than 10 µm in length.

D. Neurites in sublamina S1 had similar densities of input and output synapses in the new data set (B/K dApex2) and the subset of straight dendrites with dense synapses in the existing data set (k0725; boxed region).
