## Supplementary material for "Molecular identification of wide-field amacrine cells in mouse retina that encode stimulus orientation": Figure 6-figure supplement 2

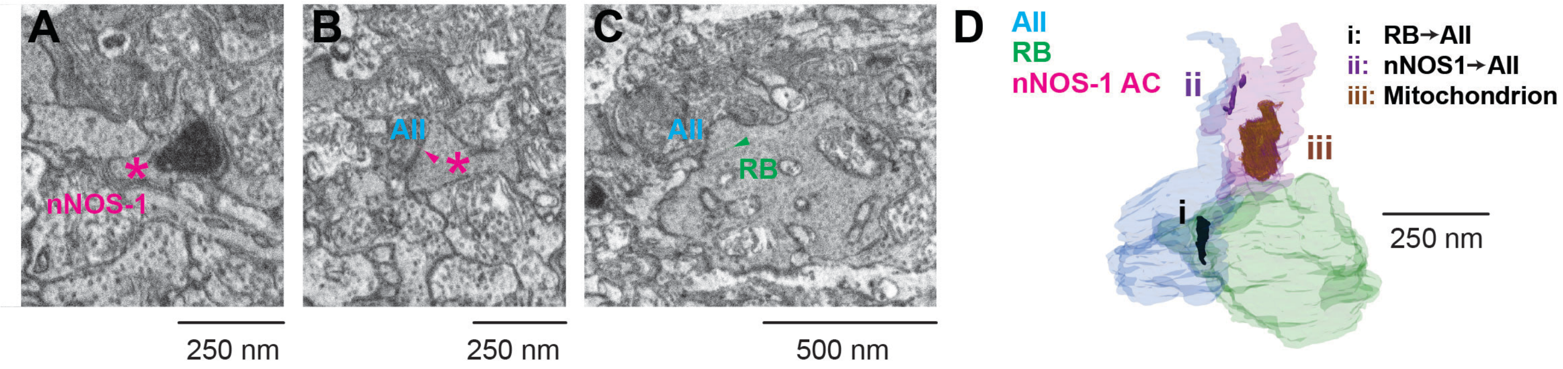


Figure 6-figure supplement 2. dAPEX2-labeled process of an nNOS-1 amacrine cell makes a synapse with an AII amacrine cell.
A. SBEM section from an nNOS-CreER;LSL-dAPEX2 retina showing a DAB+ process in the ON layer adjacent to rod bipolar cell terminals. The labeled process belongs to a presumed nNOS-1 amacrine cell.
B. Section nearby the one in A. showing a synapse from the DAB+ process to an AII amacrine cell, which was identified based on its morphology and its input from a rod bipolar cell.
C. Section nearby the one in B. showing a rod bipolar cell making a ribbon synapse with the AII amacrine cell in B.
D. 3D reconstruction showing the synapses illustrated in B. and C. and the labeled (DAB+) mitochondrion in A.
